# The Ral small GTPase is an essential regulator of Exocyst complex function in secretion

**DOI:** 10.1101/2025.08.28.672893

**Authors:** You Wu, David J. Reiner

## Abstract

Ral GTPases have long been proposed as regulators of the metazoan Exocyst, a conserved secretory vesicle-tethering complex, but direct evidence for this role has been scarce. In contrast, the well-studied yeast Exocyst relies on multiple Rab GTPases to regulate function, but yeast do not encode Ral. Using *Caenorhabditis elegans* we demonstrate that endogenous RAL-1 directly engages the Exocyst through conserved binding sites in its subunits. Loss of RAL-1 disrupts dendritic arborization of PVD sensory neurons, impairs vesicle trafficking, and causes broad developmental defects, acting both cell-autonomously in neurons and non-autonomously through supporting epithelial cells. Structure-guided genome editing of RAL-1-Exocyst interfaces produced synthetic phenotypes, underscoring the physiological importance of these contacts. Taken together, our findings establish RAL-1 as a *bona fide* regulator of the metazoan Exocyst *in vivo* and suggest that Ral-Exocyst interactions operate in parallel with other secretory pathways. More broadly, this work positions *C. elegans* as a powerful system to dissect Ral-Exocyst mechanisms across molecular, cellular, and developmental scales.

## INTRODUCTION

Ras, the founding member of the Ras superfamily of small GTPases, is the most frequently mutated human oncoprotein (Prior et al., 2020). Its best characterized oncogenic effector pathways – Raf>MEK>ERK and PI3K>PDK>AKT – are intensively studied and pharmacologically targeted (Cox et al., 2014; Stalnecker and Der, 2020). A third effector arm, mediated by guanine nucleotide exchange factors (RalGEFs) that activate the Ral subfamily of small GTPases, also contributes to tumorigenesis (Yuan et al., 2018), but its downstream mechanisms remain underexplored (reviewed in Apken and Oeckinghaus, 2021; Gentry et al., 2014; Kashatus, 2013). Mutational dysregulation of Ral has also been linked to neurodevelopmental disorders (Christen et al., 2023; Hiatt et al., 2018; Shimojima et al., 2009; Wagner et al., 2020; Xu et al., 2022).

In vertebrates, GTP-bound Ras activates four RalGEFs that in turn activate RalA and RalB, whereas in invertebrates Ras signals through a single RalGEF to a single Ral protein. Ral (<u>Ra</u>s- like) small GTPases engage binding partners distinct from other Ras family members and appear to be activated exclusively by RalGEFs (Apken and Oeckinghaus, 2021; Feig, 2003). Although RalBP1 was long considered the canonical Ral effector, genetic analyses suggest its functions are mostly independent of Ral (Cantor et al., 1995; Goldfinger and Lee, 2013; Jullien- Flores et al., 1995; Neel et al., 2012). Instead, the best studied effectors of Ral are the Exocyst subunits Sec5 and Exo84. These interactions led to the influential model that Ral regulates the Exocyst, a conserved, heterooctameric tethering complex essential for exocytosis (**Fig. 1A**). However, the underlying evidence derives mainly from data from over 20+ year-old biochemical studies: *in vitro* binding assays, yeast two-hybrid experiments, and protein fragment over- expression in cultured cells (Brymora et al., 2001; Moskalenko et al., 2002; Moskalenko et al., 2003; Sugihara et al., 2002; Wang et al., 2004).

**Figure 1.**
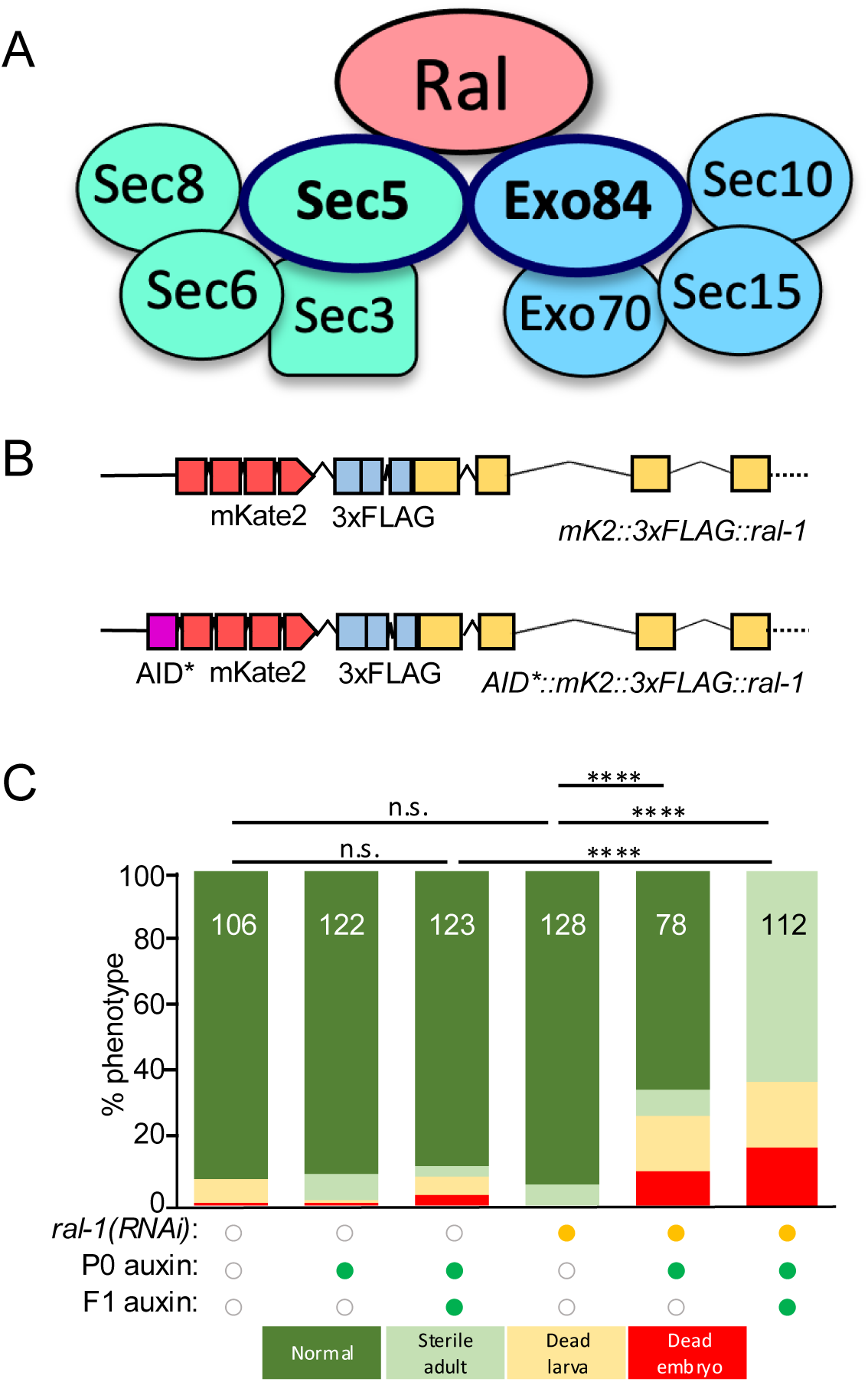
Strong depletion of RAL-1 reveals an essential role in development. **(A)** A cartoon of the central hypothesis: Ral is a key regulator of the Exocyst. See also Fig. S1A for schematics of Exocyst subunits in yeast, worms, flies and humans. **(B)** A schematic of reagents central to this study: *ral-1(re218[mKate2::3xflag::ral-1])* and *ral- 1(re218re319[AID*::mKate2::3xflag::ral-1])* (Shin *et al.,* 2018; Fakieh *et al*., 2022), the latter used for conditional depletion in all somatic cells or in specific tissues.. **(C)** Overlapping modes of AID*::RAL-1/*ral-1* depletion – auxin with somatically expressed TIR1 cofactor and *ral-1(RNAi)* – are additive as assayed by lethality. Neither *ral-1(RNAi) alone* nor auxin/TIR1- dependent depletion of AID*-tagged RAL-1 protein alone conferred growth defects, but simultaneous co-depletion conferred sterility and some lethality. A color-coded genotype key is below the graph. ****<0.0001, n.s. = not significant (Fisher’s exact test).

The Exocyst tethers post-Golgi secretory vesicles to the plasma membrane (PM), directing them to SNARE-mediated fusion sites (Hsu et al., 1996; Stalder and Gershlick, 2020; Volpiana et al., 2024). Vesicles exiting the trans-Golgi network (TGN) can be retained in the Golgi, recycled to the endoplasmic reticulum (ER), trafficked to endolysosomal compartments, or delivered to the plasma membrane (PM). Secretory routes are typically distinguished along two axes: direct vs indirect (straight from the TGN to the PM *vs.* via endosomal intermediates) and constitutive vs regulated (continuous secretion *vs.* stimulus-triggered release). Among these, the direct constitutive route is essential for delivering secreted factors, membrane proteins and newly synthesized lipid bilayer to the cell surface, and is well established to depend on the Exocyst. Whether the Exocyst also participates in indirect secretion via recycling endosomes remains unresolved.

The Exocyst belongs to the CATCHR (<u>c</u>omplexes <u>a</u>ssociated with tethering <u>c</u>ontaining <u>h</u>elical rods) group of multisubunit tethering complexes (Santana-Molina et al., 2021; Szentgyorgyi and Spang, 2023). It comprises eight subunits: Sec3, Sec5, Sec6, Sec8, Sec10, Sec15, Exo70, and Exo84 (EXOC1-8 in mammals; **Fig. S1A**). In general, “Sec” subunits are essential for secretion, whereas Exo70 and Exo84 are partially dispensable. The Exocyst is essential in yeast and mammals (Schekman, 1992; Wang et al., 2015). In contrast, functional analysis in metazoans is complicated by maternal loading: maternal-plus, zygotic-minus (M+Z-) of most Exocyst subunits in *Drosophila* can complete embryogenesis, and, in the case of the developmentally simpler *C. elegans* with far fewer cells, reach sterile adulthood (Armenti et al., 2014a; Hyenne et al., 2015; Murthy et al., 2003; Murthy et al., 2005; Murthy and Schwarz, 2004). This maternal rescue obscures essential early roles and mechanisms of *in vivo* regulation.

Cryo-EM studies of yeast Exocyst suggest that subunits assemble into two heterotetrameric subcomplexes (S1 and S2) via multiplexed “CorEx” coiled-coil interfaces. S1 and S2 then interdigitate to form the heterooctamer (**Fig. S1B-E**; Mei et al., 2018). Other work depicts the Exocyst as more extended and flexible, consistent with earlier EM or yeast subunit tethering and tagging experiments (Hsu et al., 1998; Picco et al., 2017). These discrepancies – between yeast and mammalian systems, *in vitro* and *in vivo* contexts, and cross-linked *vs.* native preparations – underscore that the architecture of the complex *in vivo* remains intriguing but unresolved.

Small GTPases are major regulators of the Exocyst in yeast: Rabs, Rho, and Cdc42 help specify vesicle identity and spatial targeting (Wu and Guo, 2015; Wu et al., 2008). Ral, absent in yeasts but present in metazoans, has been proposed to regulate Exocyst assembly or function through Sec5 and Exo84. Yet this model remains largely circumstantial, with no demonstration of endogenous Ral–Exocyst interactions in living animals.

In *C. elegans*, *ral-1* deletion homozygotes from heterozygous mothers (M+Z-) progress to sterile adults, mirroring the phenotypes of Exocyst deletions (Armenti et al., 2014a; Hyenne et al., 2015; Shin et al., 2019; Zand et al., 2011). Deletion studies in *Drosophila* and mice complicate the picture but do not establish a direct role for Ral in Exocyst regulation (Balakireva et al., 2006; Peschard et al., 2012). Moreover, though extensively used, dominant-negative GTPase mutants are susceptible to artifacts and misinterpretation (Feig, 1999).

Thus, despite two decades of widespread citation, the idea that Ral is an obligate regulator of the Exocyst in metazoans still lacks definitive *in vivo* support. This uncertainty complicates interpretation of the role of Ral in Ras-driven oncogenesis: if Ral governs secretion, its contribution to tumorigenesis could reflect disrupted vesicle trafficking rather than conventional signal transduction. In contrast, signaling roles of RalGEF>Ral are firmly established: in *C. elegans*, Ras>RalGEF>Ral promotes secondary VPC fate, and RalGEF>Ral acts permissively in cell and neurite migrations where the Exocyst is dispensable (Mardick et al., 2021; Shin et al., 2019; Shin et al., 2018; Zand et al., 2011).

To address this longstanding question, we tested whether Ral is required for Exocyst-dependent exocytosis in a metazoan. Using *C. elegans*, we combined targeted genetic perturbations, conditional protein depletion, and high-resolution phenotypic analyses to test whether RAL-1 is required for Exocyst function. We further examined whether endogenous RAL- 1 physically interacts with and co-localizes with endogenous Exocyst subunits *in vivo*. Finally, we analyzed structure-guided point mutations in two predicted Ral-Exocyst interfaces and found that each impaired Exocyst function. Taken together, our findings demonstrate that RAL-1 is essential for Exocyst-dependent secretion and physically associates with the Exocyst *in vivo*. Rather than serving as an obligate structural component, RAL-1 likely acts as a regulatory cofactor, analogous to GTPase modulators of other multisubunit tethering complexes. This establishes a direct link between Ral and the Exocyst in an intact animal and provides a tractable system to dissect their roles in secretion, signaling, and development.

## RESULTS

### *C. elegans* Ral is required for development and fertility

We previously tagged the endogenous RAL-1A isoform at the N-terminus with the red fluorescent protein mKate2 (also known as mK2) and 3xFLAG epitope (Shin et al., 2018). To enable conditional depletion, we inserted sequences encoding the minimal 44 amino acid auxin-inducible degron (AID*) at the 5’ end of this modified *ral-1* gene (**Fig. 1B**; Fakieh et al., 2022).

We then generated animals expressing AID*::mK2::3xFLAG::RAL-1 from the endogenous locus, along with somatically expressed TIR1, the F-BOX substrate recognition subunit of the SCF E3 ubiquitin ligase, required or auxin-dependent degradation (Zhang et al., 2015).

Anti-FLAG immunoblotting confirmed that degradation of AID*::mK2::3xFLAG::RAL-1 was dependent on both TIR1 expression and auxin treatment (**Fig. S1F**). Depletion occurred rapidly but remained incomplete, and increasing auxin concentration did not improve depletion efficiency (**Fig. S1G**). Under these conditions, animals appeared superficially wild type. In a previous study, injected *ral-1* dsRNA or bacterially mediated *ral-1*-directed RNAi in an *rrf-3* RNAi hyposensitive animal failed to cause any growth defects; only a signaling phenotype was observed (Zand et al., 2011). Taken together, these results suggest the threshold for RAL-1 in Exocyst function may not require much protein.

Single perturbations targeting *ral-1* – either two generations of auxin-dependent depletion or one generation of *ral-1(RNAi)* via bacterial feeding – did not cause strong phenotypes (**Fig. 1C**). However, combining one generation of auxin with concurrent *ral-1(RNAi)* produced mild growth defects. Only the combination of two generations of auxin with concurrent *ral-1(RNAi)* led to complete sterility and low penetrance larval and embryonic lethality (**Fig. 1C**, column 4 *vs.* 5 *vs.* 6). These phenotypes closely resemble those reported for the *ral-1* deletion alleles *tm2760* and *tm5205* (Hyenne et al., 2015; Shin et al., 2019; Zand et al., 2011). Notably, prior studies using ZIF-1/ZF1-mediated depletion of RAL-1 produced even stronger phenotypes (Armenti et al., 2014a; Armenti et al., 2014b), suggesting that auxin + RNAi results in partial loss of function, a deduction further supported by **Fig. 2C**.

**Figure 2.**
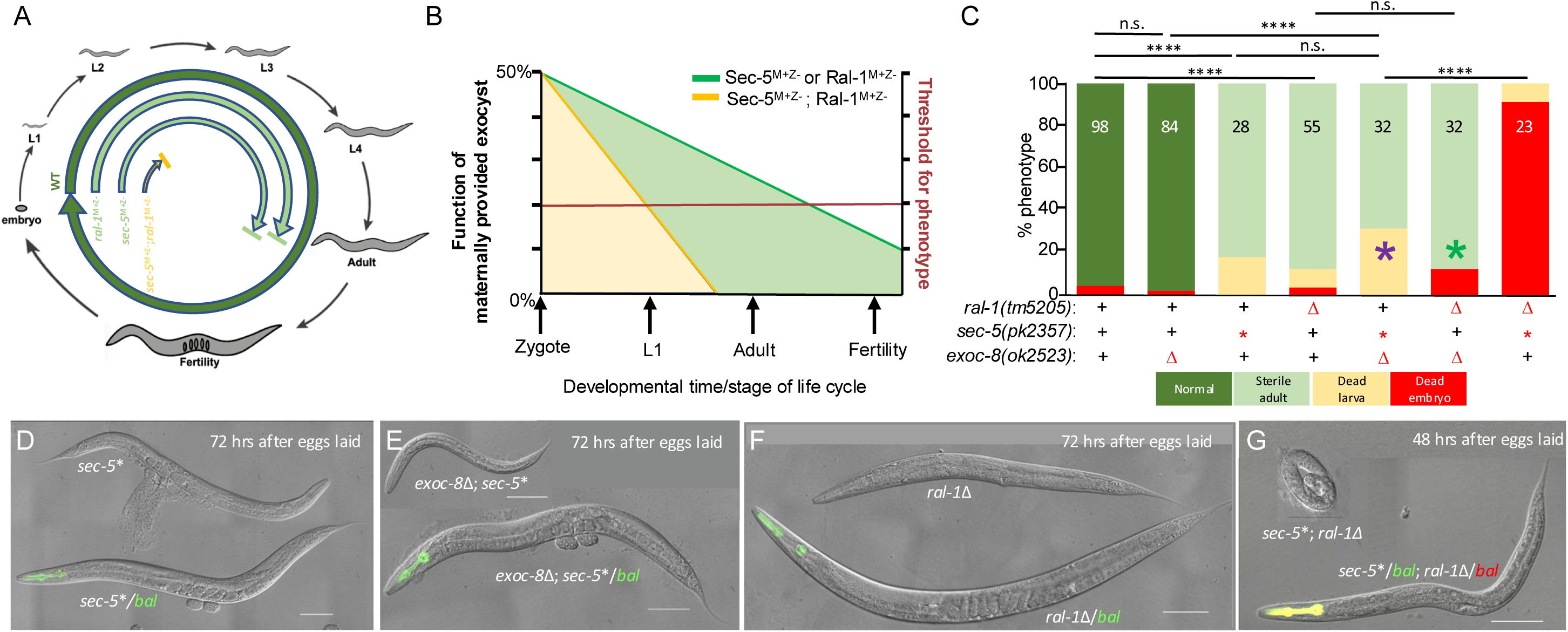
Reduction of *ral-1* function aggravates growth defects of deficient exocyst mutants. **(A)** The life cycle of hypothetical M+Z- single mutant vs. double mutant animals. Wild type (WT; dark green) completes the life cycle, M+Z- sec-5* or *ral-1*Δ animals (light green) have sufficient maternal Ral and exocyst gene product to reach adulthood but not fertility. M+Z- *sec- 5*+ral-1*Δ double mutant animals (yellow) have insufficient gene product to support full development. **(B)** A hypothetical graph of the decrease in maternally provided gene product over developmental time and the functional requirements of gene product in single vs. double mutants. M+Z- animals likely start with 50% functional gene product that decreases over time. The X-axis indicates developmental stages while the red line and Y-axis label on the right indicate a hypothetical threshold of gene product function required to support continued development. The green line denotes decrease in single mutant M+Z- gene product over time, the yellow line the expected decrease in double mutant M+Z- gene product over time. Due to maternal rescue, neither mutant is null for function. **(C)** Graphed developmental progression of single and double mutants by larval stage, color key is below. Purple asterisk = *exoc-8*; *sec-5* double mutants did not arrest or die at an earlier stage, but animals were much smaller than single mutants (**Fig. S2A-D**). Green asterisk = *exoc-8*; *ral-1* double mutants did not arrest or die at an earlier stage, but animals were much heterozygote segregates only. Small number of double homozygotes. **(D-G)** Representative DIC images of control single vs. double mutant animals. Genetic balancers for (**D**) *sec-5* (green pharynx) and (**E**) ral-1 (red pharynx) were *mIn1[myo-2p>gfp]* and *qC1[myo-2p>rfp]*, respectively. The pharynx of the double balancer animal is shown as yellow. bal = genetic balancer, Δ = deletion, * = nonsense mutation. White scale bar = 100 mm; black = 50 mm. Genotype key is below each graph. ****<0.0001, n.s. = not significant (Fisher’s exact test).

Importantly, these combined depletion phenotypes validate that the M+Z- phenotypes (maternally rescued, see below) observed with deletion alleles reflect the true hypomorphic loss of *ral-1* activity. This conclusion is critical because knockout alleles originate from heavily mutagenized backgrounds where phenotypes may be confounded despite repeated outcrossing. Thus, conditional RAL-1 depletion offers a robust and interpretable tool for dissecting Ral function.

*ral-1* encodes two isoforms. Exon 1a encodes the evolutionarily conserved N-terminus of RAL-1A, the canonical ortholog of metazoan Ral GTPases (**Fig. S1H**; Zand et al., 2011). Exon 1b, located several hundred bp upstream of the ATG initiator methionine codon of exon 1a, encodes a longer, N-terminal extension not conserved in other metazoans but found in all nematode species examined. By RNAseq, exon 1b expression is ≤13% of total *ral-1* expression, with exon 1a comprising the rest.

To eliminate exon 1b expression, we used CRISPR/Cas9-mediated STOP-IN gene disruption (Wang et al., 2018) to insert a 43 bp disruption cassette into exon 1b of the *ral- 1(re218[mK2::3xFLAG::ral-1])* background, generating two independent alleles, *re470* and *re471* (**Fig. S1I**). Both mutants are viable and fertile with no discernable change in FLAG-tagged protein levels (**Fig. S1J**). *re471* did not reduce Exocyst function (**Fig. S1K**) using assays established in this study (see below). *re471* did not reduce signaling function of *ral-1* in a standard assay established in our lab *re471* (**Fig. S1L**; Shin et al., 2019; Shin et al., 2018; Zand et al., 2011). These findings suggest that the mRNA and protein species encoded by exon 1b is a nematode-specific innovation that is dispensable for canonical Ral functions. We therefore do not discuss exon 1b further in this study.

### Genetic interactions among RAL-1 and Exocyst components support shared roles in larval development

*C. elegans* RAL-1, SEC-5 and all SEC-named Exocyst components are essential proteins; homozygous deletion mutants cannot be propagated as stable lines. To study their function, we analyzed animals derived from heterozygous mothers, referred to as maternal-plus, zygotic- minus (M+Z-) genotypes. In this context, homozygous mutant zygotes from heterozygous mothers are expected to inherit approximately 50% of the maternal wild-type RNA and protein pool (**Fig. 2A**). In contrast, EXOC-8 and EXOC-7, like their yeast orthologs Exo84 and Exo70, are non-essential for viability, though they still contribute to Exocyst function.

We observed that *ral-1(tm5205)* M+Z- mutants grow to adulthood and are sterile, while *sec- 5(pk2357)* M+Z- mutants also reach adulthood but frequently rupture at the vulva and produce a handful of inviable embryos (**Fig. 2C,D,F**). Previous work using ZIF-1/ZF1 degron-mediated depletion to generate maternal-minus, zygotic minus (M-Z-) mutants of RAL-1, SEC-5, and SEC- 8 reported 100% lethality at the first larval stage (L1), along with some embryonic death (Armenti et al., 2014a). This effect was attributed to severe disruption of the polarized epithelial canal of the excretory cell, a structure critical for osmoregulation and analogous to the mammalian kidney.

However, because null mutant in Exocyst components arrest development entirely, they are not well suited for analyzing the functional contributions of RAL-1 to the Exocyst. Classical genetic pathway analysis ideally uses null mutations to assign linear relationships (Avery and Wasserman, 1992; Hereford and Hartwell, 1974), but genetic interaction studies probing cooperation within shared processes are more informative under sensitized conditions, such as partial loss-of-function alleles, which can reveal synthetic effects (Simon et al., 1991; Simon et al., 1993). M+Z- animals are such partial loss-of-function mutants. Yet maternal rescue is an unusual form of loss-of-function condition, where degree of reduced function increases over time as maternal product decreases via decay or dilution with animal growth. We illustrate the theoretical contributions of parallel maternally provided gene products in **Fig. 2A** and their theoretical impact on development in **Fig. 2B**. In other words, M+Z- deficient RAL-1 function is likely to exacerbate M+Z- deficient Exocyst phenotypes but not strong or null Exocyst phenotypes: development fails when SEC-5 or RAL-1 gene products fall below certain threshold levels, illustrated in hypothetical schematics (**Fig. 2A,B**).

We therefore used a panel of mutants, mostly from the *C. elegans* gene knockout consortium, to assess genetic interactions between *ral-1* and its putative Exocyst binding partners *sec-5* (Sec5) and *exoc-8* (Exo84). As a starting point, the *exoc-8(ok2523)* deletion of a non-essential subunit was crossed with the nonsense allele *sec-5(pk2357)* (Frische et al., 2007). This double mutant was much more developmentally impaired than either single mutant (**Fig. 2C**,**E**), producing viable but markedly smaller animals (**Fig. S2A-D**). This interaction was replicated with second alleles: *exoc-8(ok2523); sec-5(tm1443)*, and *exoc-7(ok2006); sec-5(pk2357)*, both of which showed enhanced phenotypes compared to single mutants (**Fig. S2E,F**).

Crucially, the *sec-5(pk2357)*; *ral-1(tm5205)* M+Z- double mutant exhibited significantly more severe defects than either single mutant, with most animals dying as embryos (**Fig. 1C,G**). This finding was also recapitulated with second alleles including *ral-1(tm2706); sec-5(tm1443)* and *ral-1(tm5205); exoc-7(ok2006)*, both of which showed enhanced phenotypes compared to single mutants (**Fig. S2E,F**). Not all double mutant combinations caused arrest. For example, *ral- 1(tm5205)* or *ral-1(tm2706)* double mutants with *exoc-8(ok2523)* did not arrest but exhibited decreased body length and width (**Fig. S2G,H**), perhaps indicative of less severe Exocyst dysfunction. By contrast, *ral-1(tm5205); exoc-7(ok2006)* animals arrested as larvae (**Fig. S2F**). These differences may reflect variation in perdurance of gene product (mRNA, protein) of each subunit combined with variance in contribution of each subunit. Also, the mechanism of developmental defect here is immaterial: we are using developmental progression and animal growth as a means to study genetic interactions, not the end in itself. (We speculate that defective excretory system is causing these growth deficits, as studied previously (Armenti et al., 2014a).

Taken together, these genetic interactions between partial loss-of-function alleles of *ral-1* and components of the Exocyst support a model in which RAL-1 is a critical regulator of Exocyst function during development. However, these interactions could, in principle, also result from parallel roles in distinct but converging secretory pathways within the endomembrane system. Therefore, while genetic evidence supports a collaborative role, we pursued several complementary mechanistic studies to directly assess the functional relationship between RAL- 1 and the Exocyst.

### RAL-1 is required for proper exocyst-dependent PVD dendritic arborization

During larval development, the bilateral mechanosensory PVD neurons extend an elaborate and stereotyped network of dendrites, a process termed “arborization,” which enables them to tile much of the animal’s body. Due to this extensive branching and its reproducible morphology, the PVD neuron serves as an established model for analyzing the functions of trafficking proteins, including components of the Exocyst complex and regulators of recycling endosomes such as RAB-10 (Chen et al., 2006; Shi et al., 2010). Arborization is readily visualized using a PVD-specific GFP reporter that labels the full dendritic architecture in staged larvae (Taylor et al., 2015; Zou et al., 2015).

To examine RAL-1 localization *in vivo*, we used the *ral-1(re218[mK2::3xFLAG::ral-1])* knock-in strain (Shin et al., 2018) that expresses a tagged, red-fluorescent endogenous RAL-1 fusion protein. mK2::RAL-1 red fluorescent signal was detected in the PVD cell body, consistent with endogenous RAL-1 expression in these neurons (**Fig. S3A**).

The PVD dendritic tree is organized into four stereotyped layers: primary (1°), secondary (2°), tertiary (3°), and quaternary (4°) branches (**Fig. S3B**). Consistent with prior studies (Taylor et al., 2015; Zou et al., 2015), we observed reduced dendritic complexity in *exoc-8* deletion or in *sec- 5(pk2357)* M+Z⁻ mutants (**Fig. 3A,B,E; Fig. S3C,D**). Similarly, *ral-1(tm5205)* M+Z- mutants exhibited significantly diminished PVD arborization (**Fig. 3A,C,E**), as did animals depleted of RAL-1 via auxin-inducible degradation (AID*) in somatic tissues (**Fig. 3F**; **Fig. S3E,F**). These phenotypes were qualitatively like those observed in *exoc-8* and *sec-5* mutants, suggesting a shared requirement for RAL-1 and the Exocyst in supporting PVD arborization.

**Figure 3:**
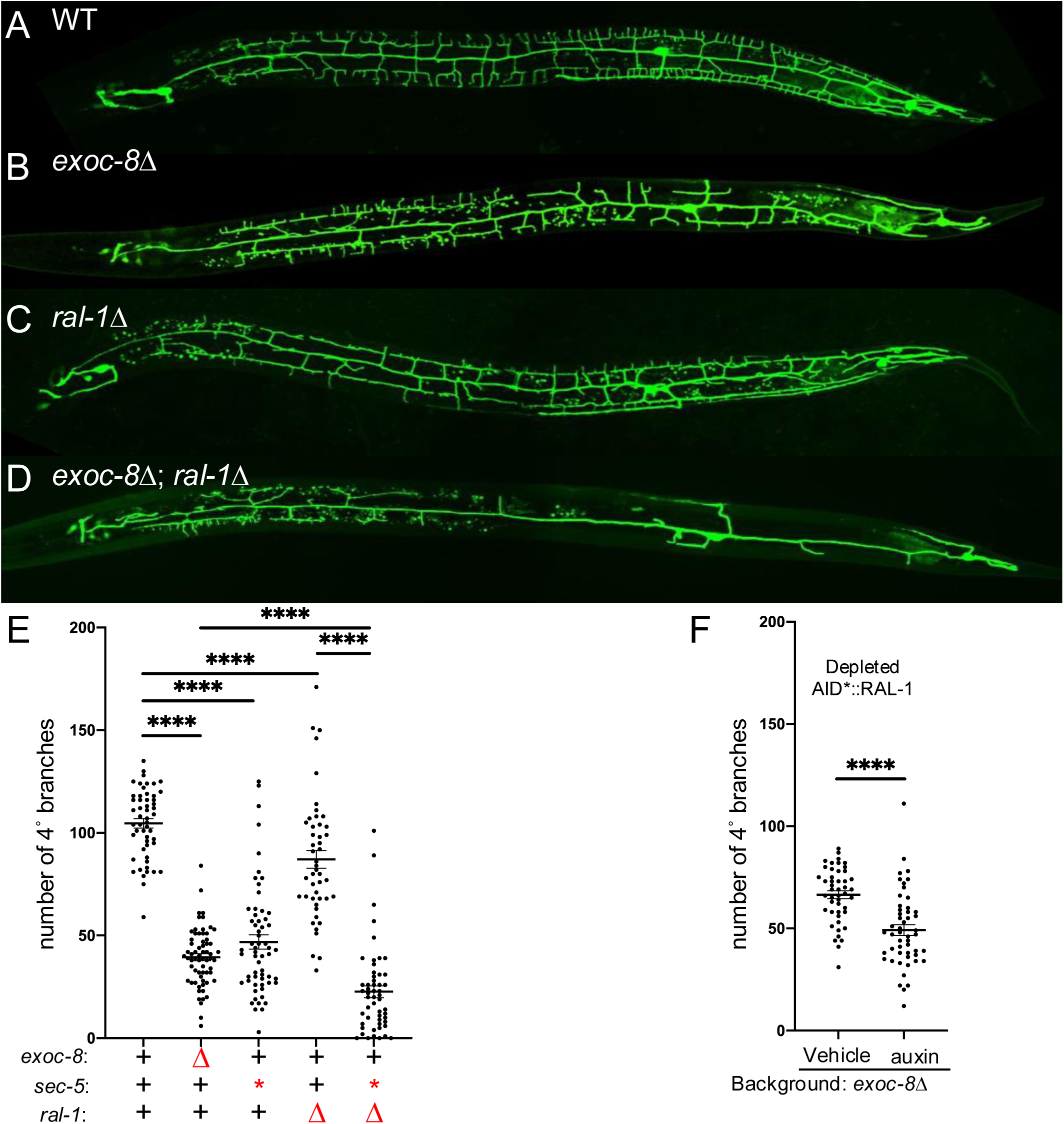
RAL-1 is required for full exocyst-dependent PVD dendritic arborization. A-D) Maximum intensity projection of z-series confocal photomicrographs of GFP-labeled PVD mechanosensory neurons (*wdIs52[ser-2(prom3)p>gfp])*) of (**A**) wild type, (**B**) *exoc-8*(*ok2523*), (**C**) *ral-1*(*tm5205*) M+Z-, and **D)** *exoc-8*(*ok2523)*; *ral-1*(*tm5205)* M+Z- double mutant animals reveal strong genetic interactions among these reduction of exocyst mutants. Anterior is left for all four animals. **E)** The *exoc- 8(ok2523)*; *ral-1(tm5205)* double mutant exhibits more severe defects in 4° dendritic branching than each single mutant. **F)** Animals of genotype *ieSi57[eft-3p>TIR1::mRuby]*; *ral-1(re218re319[AID*::mKate2::3xFLAG::ral-1])* for ubiquitous TIR1 expression display auxin-dependent enhancement of 4° dendritic branching defect conferred by *exoc-8*(*ok2523*). ****<0.0001 (*t*-test)

Strikingly, the *exoc-8(ok2523); ral-1(tm5205*) M+Z- double mutant displayed a much more severe dendritic arborization defect than either single mutant animal (**Fig. 3B-E; Fig. S3C,D**). We were unable to assess dendritic arborization in *sec-5*; *ral-1* M+Z- double mutants, as these animals did not survive to the late L4 larval stage when PVD development is normally evaluated (**Fig. 2C**). Taken together, these data support a role for RAL-1 as a key upstream regulator of Exocyst-dependent neuronal morphogenesis.

### RAL-1 is required for proper Exocyst-dependent vesicle trafficking in PVD dendrites

We used DMA-1::GFP, a GFP-tagged integral membrane protein that functions as a dendritic guidance receptor in PVD neurons (Ziegenfuss and Grueber, 2013), to monitor trafficking and exocytosis of vesicle cargo. Defects in exocytosis result in increased numbers of DMA-1::GFP punctae in the primary (1°) dendrites and reduced fluorescence intensity of these vesicles (**Fig. 4A**; see Methods; (Taylor et al., 2015; Zou et al., 2015).

**Figure 4:**
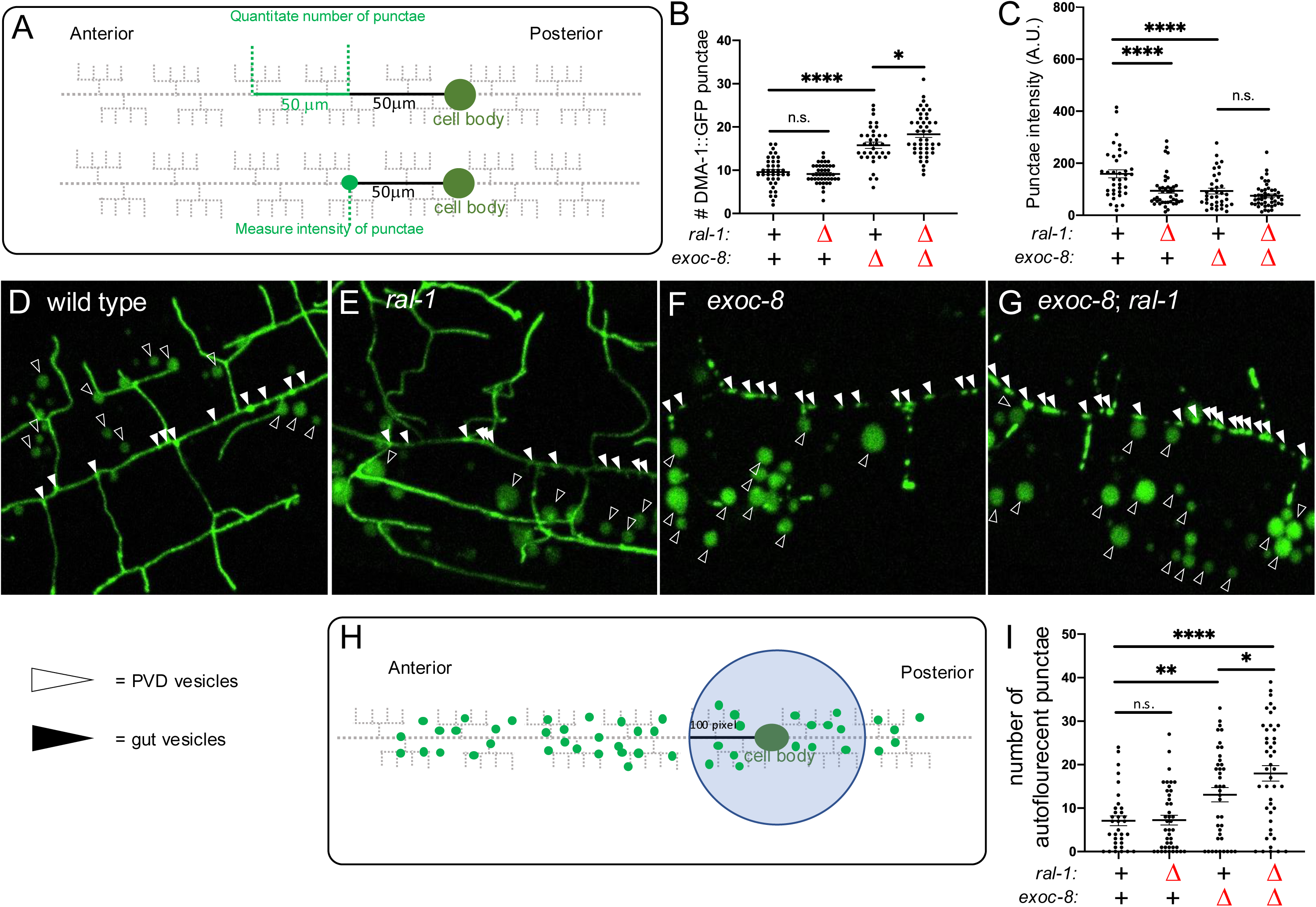
RAL-1 is required for proper exocyst-dependent vesicle biogenesis and trafficking in PVD mechanosensory neurons. **(A)** A schematic of the method for quantitating number and intensity of punctae of DMA::GFP vesicle cargo (*qyIs368[ser-2prom3p>dma-1::gfp]*) in 1° dendrites of PVDs. Defective exocyst complex increases number but decreases intensity of vesicles in the 1° dendrite of PVDs (Taylor *et al*., 2015; Zou *et al*., 2015). **(B)** M+Z- RAL-1 did not increase the number of proximal vesicles but mildly enhanced the defect of *exoc-8(ok2523).* **C)** M+Z- *ral-1* decreased intensity of DMA- 1::GFP punctae as a single mutant but did not aggravate the decreased intensity of the *exoc-8(ok2523*) mutant. **(D-G)** Confocal photomicrographs of the *qyIs368[ser-2prom3p>DMA-1::GFP]* vesicular cargo protein (a transmembrane patterning receptor) in secretory vesicles in PVDs. Vesicles bearing DMA-1::GFP cargo are indicated by solid arrowheads. Hollow arrowheads indicate autofluorescence from Lysosome-related organelles (LROs; Soukas *et al*., 2012) in the intestine, which is captured by the confocal z-stack imaging the PVDs and we noticed were altered by mutation of *exoc-8* and *ral-1*. **(H)** A schematic of the method for quantitating number of autofluorescent intestinal vesicles in a 100 pixel diameter circle area around the cell body. (**I)** *exoc-8* significantly increased the number of autofluorescent punctae. *exoc-8*; *ral-1* has significantly more punctae than *exoc-8*. *<0.05, **<0.01, ***<0.001, ****<0.0001, n.s. = not significant (*t-*test).

As expected, deletion of *exoc-8* significantly increased number of vesicles (**Fig. 4B,D,F**; white arrowheads). In contrast, M+Z- *ral-1* deletion did not increase vesicle number (**Fig. 4B,D,E**). However, the *exoc-*8; *ral-1* double mutant showed a further increase in vesicle number compared to the *exoc-8* single mutant (**Fig. 4B,F,G**). Both *exoc-8* deletion and *ral-1* M+Z- deletion decreased vesicle intensity (**Fig. 4C-F**), but this reduction was not observed in the double mutant (**Fig. 4C-G**). These data are consistent with RAL-1 promoting vesicle exocytosis and supports a role for RAL-1 as a regulator of the Exocyst.

We also examined Exocyst and RAL-1 function in the intestine, where autofluorescent vesicle-like lipid storage droplets serve as an additional readout for vesicle trafficking. We quantified the number of autofluorescent punctae within a 100 pixel-diameter region centered on the PVD cell body (**Fig. 4H**) in the shown maximal intensity projection of the z-stacks. *exoc- 8* deletion, but not *ral-1* M+Z- deletion, significantly increased number of punctae compared to wild type (**Fig. 4D-F,I**; black arrowheads). However, combining *ral-1* M+Z- with *exoc-8* deletion led to a further increase in autofluorescent punctae (**Fig. 4E-G,I**). These findings, in both PVD dendrites and the intestine, suggest that RAL-1 acts with the Exocyst to promote exocytosis of secretory cargos. The PVD findings suggest that RAL-1 and the Exocyst complex are required not only for vesicle tethering at the target membrane, but also for earlier steps in vesicle assembly, transport, and secretion.

### RAL-1 functions both cell-autonomously and non-autonomously

Because the Exocyst complex mediates direct exocytosis from the Golgi, we hypothesized that both the Exocyst and RAL-1 function non-autonomously – acting in both migrating PVD dendrites and the underlying epithelial substrate – to facilitate the secretion of factors required for PVD dendrite morphogenesis (Inberg et al., 2019; Sundararajan et al., 2019; Yang and Chien, 2019). To test this hypothesis, we used tissue-specific chemical-genetic depletion of endogenous AID*::mKate2::RAL-1 using the auxin-inducible degradation (AID*) system (Fakieh et al., 2022; Zhang et al., 2015; **Fig. 1B**). Tissue-specific degradation was achieved using previously validated neuron- or epithelial-specific (termed the hypodermis in *C. elegans*) expression of the TIR1 F-box protein combined with a stoichiometric internal control, TagBFP2::AID*::NLS, which labels the nuclei of auxin-responsive cells (blue nuclei; see Methods; (Ashley et al., 2021).

To enhance sensitivity to RAL-1 depletion, animals in the arborization assays were mutant for *exoc-8* and harbored a PVD-specific GFP marker. For the DMA-1::GFP cargo trafficking experiments, animals were *exoc-8(*+), ensuring that the observed phenotypes were attributable solely to depletion of RAL-1.

As expected, neuronal-specific depletion of RAL-1 in *exoc-8* mutants caused severe PVD arborization defects relative to vehicle control (**Fig 5A-F**). In *exoc-8(+)* animals, neuronal depletion of RAL-1 resulted in increased number and reduced intensity of DMA-1::GFP-positive vesicles, consistent with trafficking defects (**Fig. 6A-D’’**). Similarly, epithelial-specific depletion of RAL-1 in *exoc-8* mutants caused pronounced PVD arborization defects (**Fig 5G-L**), and in *exoc-8(+)* animals, increased number and decreased intensity of DMA-1::GFP-positive vesicles (**Fig. 6E-H’**). In both tissues, TagBFP2::AID*::NLS nuclear signal was abolished by auxin treatment, confirming effective depletion (**Fig. 5B** *vs.* **5C**; **Fig. 6C-C’’** *vs.* **6D-D”**; **Fig. 5H** *vs.* **5I**; **Fig. 6G,G’** *vs.* **6H,H’**).

**Figure 5:**
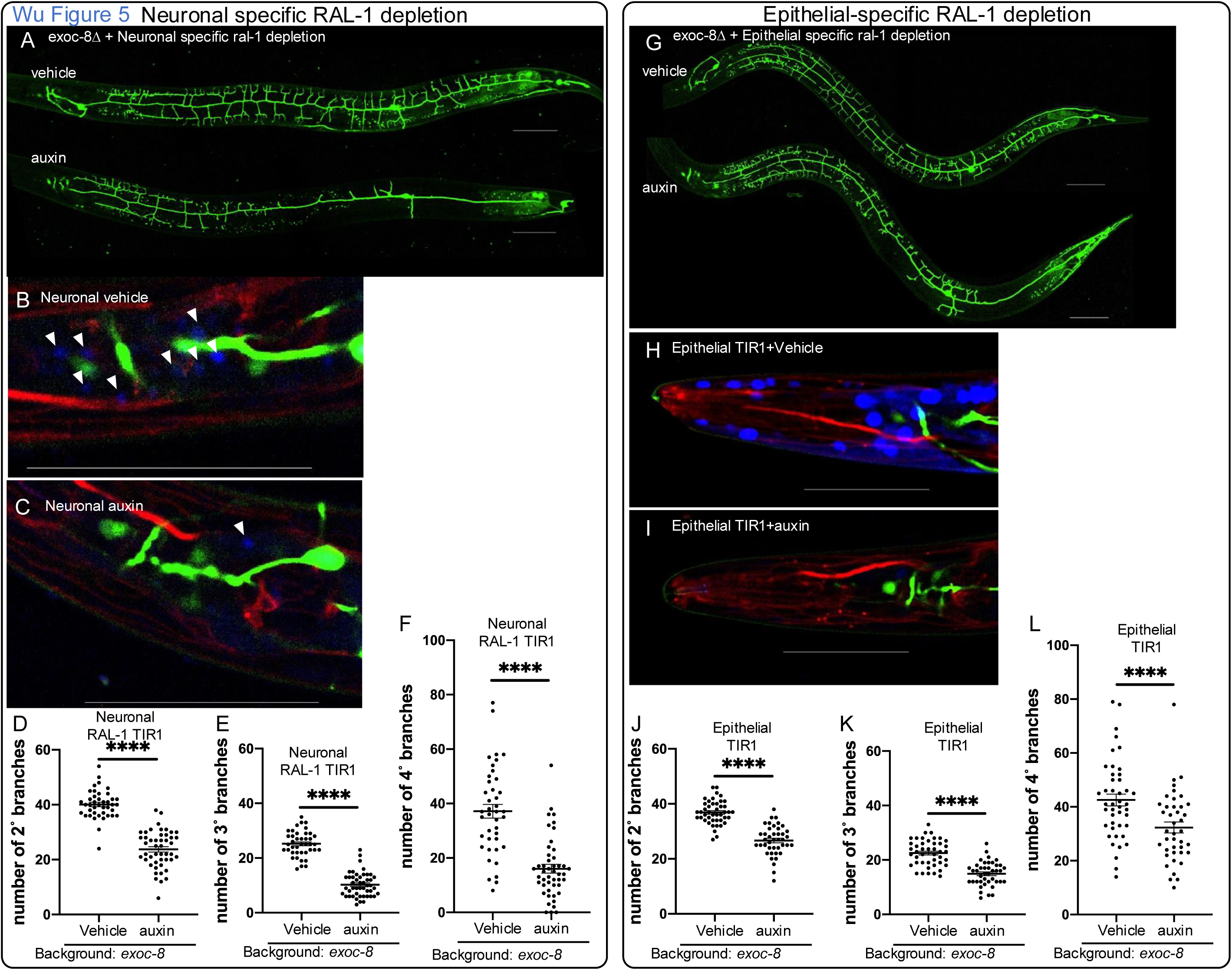
RAL-1 functions both cell autonomously and non-autonomously to regulate PVD dendrite arborization. (A-. **C)** Maximum intensity projection of z-series of confocal photomicrographs and quantification of *exoc-8(ok2523)*; *ral- 1(re218re319[AID*::mK2::3xFLAG::RAL-1])* animals, plus the *reSi7[rgef-1p>TIR1::F2A::mTagBFP2::AID*::NLS]* transgene for neuronal-specific expression of TIR1. Anterior is left for all animals. Neuronal-specific depletion of RAL-1 resulted in substantial decrease of 2° **(D)**, 3° **(E)** and 4° **(F)** dendritic branches of PVDs (*wdIs52[ser-2(prom3)p>gfp])*) in the *exoc- 8(ok2523*Δ) background. **(A)** The upper and lower animals were treated with vehicle and auxin, respectively. **(B)** A three-color zoom of the head of a vehicle-treated animal. Red = endogenous RAL-1, green = PVD marker, blue = BFP::AID*::NLS internal positive control for efficacy of auxin and neuronal-specific expression of TIR1 co-factor. Arrowheads indicate blue neuronal nuclei as internal controls for auxin efficacy. **(C)** A three-color zoom of the head of an auxin-treated animal reveals depletion of BFP::AID*::NLS internal control. The arrowhead indicates a single weak neuronal signal; others are depleted. **(G-I)** The same PVD experiments as in (**A-C**) but with epithelial-specific expression of TIR1 from the *reSi2[col-10- 1p>TIR1::F2A::mTagBFP2::AID*::NLS]* transgene, vehicle vs. auxin. **(J-L)** Epithelial-specific depletion of RAL-1 results in substantial decrease of 2° **(J)**, 3° **(K)** and 4° **(L)** dendritic branches of PVDs (*wdIs52[ser-2(prom3)p>gfp])*) in the *exoc- 8(ok2523)* background. Scale bars = 50 µm. **** <0.0001 via *t-*test.

**Figure 6.**
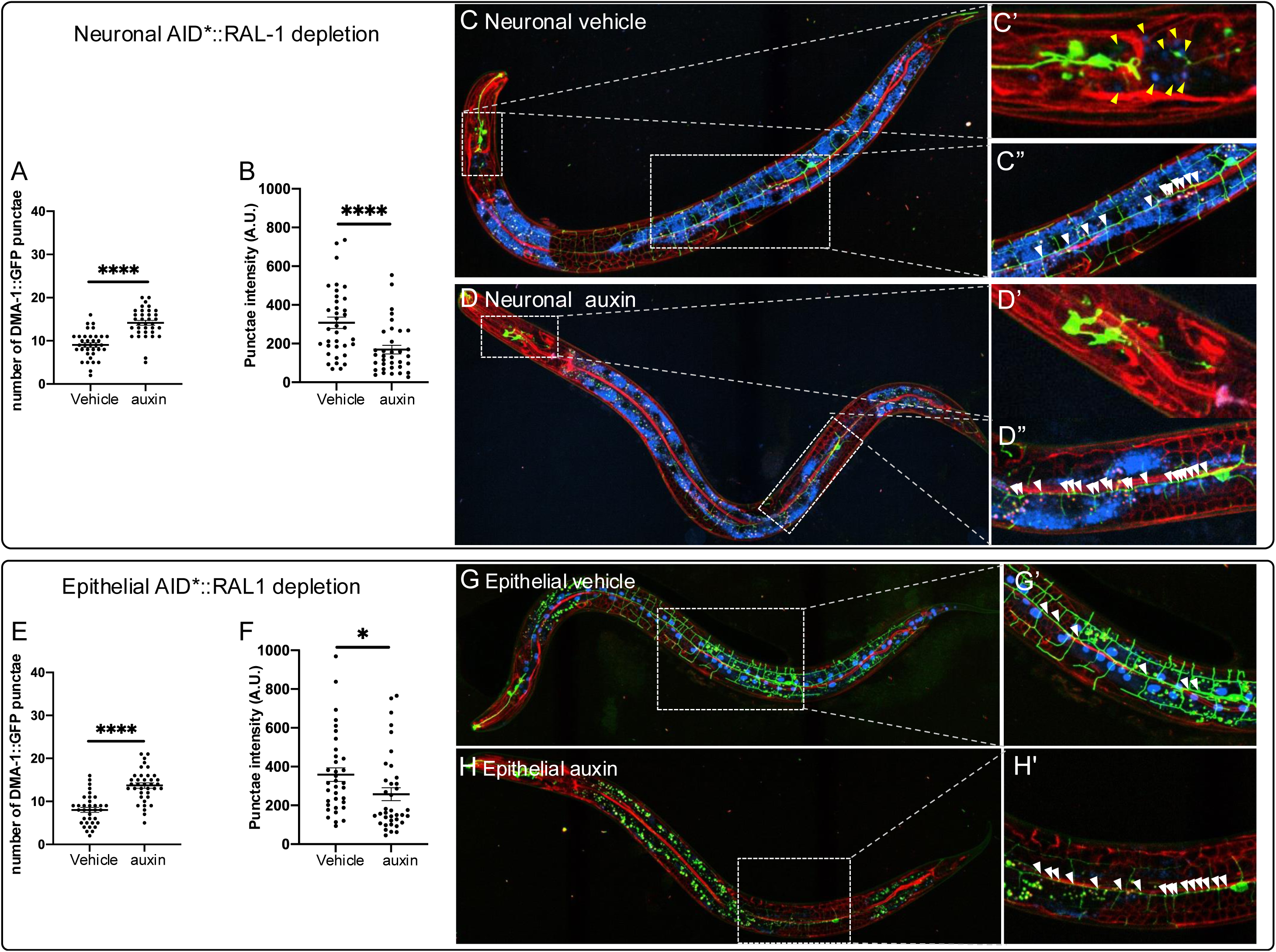
RAL-1 functions cell autonomously and non-autonomously to contribute to vesical trafficking in PVDs. (A,B) Neuronal-specific depletion of AID*::RAL-1 resulted in increased retention **(A)** and decreased intensity of DMA-1::GFP-containing vesicles **(B),** characteristic of decreased exocyst function. **C-C’’)** Confocal photomicrograph and zoom views of AID*::RAL-1 with neuronal-specific expression of TIR1 on vehicle control. **D-D’’)** Confocal photomicrograph and zoom views of AID*::RAL-1 with neuronal-specific expression of TIR1 grown on auxin for neuronal-specific depletion of AID*::RAL-1. Diffuse blue signal is intestinal autofluorescence in the 405 nm channel, required because of the long exposure time to visualize the faint BFP::AID*::NLS control in neurons. **E,F)** Epithelial-specific depletion of AID*::RAL-1 resulted in increased retention and decreased intensity of DMA- 1::GFP-containing vesicles characteristic of decreased exocyst function **G-G’’)** Photomicrograph and zoom views of vehicle control animal with hypodermal-specific depletion of AID*::RAL-1. Arrowheads indicate DMA-1::GFP-positive vesicles. **H-H’’.** Image and zoom-in views of treated animal of hypodermal-specific AID*::RAL-1 depletion. **C,D,G,H)** Red = endogenous RAL-1, green = DMA-1::GFP-containing vesicles in PVDs, blue = BFP::AID*::NLS internal positive control. Yellow arrowheads point to un- depleted blue nuclei of internal control for tissue-specific TIR1. White arrowheads indicate DMA-1::GFP-positive secretory vesicles. White arrowheads indicate DMA-1::GFP-positive vesicles. Blue intestinal autofluorescence is not visible because epithelial BFP::AID*::NLS is much brighter than neuronal BFP::AID*::NLS, and hence only requires short exposure time. *<0.05, ****<0.0001 via *t-*test.

Interestingly, tissue-specific depletion of RAL-1 produced stronger effects than observed in M+Z- *ral-1* mutants (**Fig. 3,4**). We speculate that auxin-induced depletion is consistent throughout larval development, thus reducing RAL-1 levels during critical windows of PVD outgrowth. In contrast, M+Z- loss of *ral-1* likely allows higher early expression from maternally loaded transcripts and/or protein, with gradual decline over development, potentially sparing earlier stages of elaborating dendritic structure.

In this case, differences in TIR1 expression levels are unlikely to explain the observed differences. Indeed, neuronal TIR1 expression (behind the *rgef-1* promoter) is relatively weak, while epithelial TIR1 expression (behind the *col-10* promoter) is strong, as confirmed by TagBFP2::AID*::NLS expression levels (our data; Ashley et al., 2021).

Together, these findings demonstrate that RAL-1 acts both autonomously within neurons to support dendritic outgrowth and non-autonomously within the epithelium to mediate secretion of factors that shape PVD architecture.

### Endogenous RAL-1 co-localizes with endogenous SEC-5 and EXOC-8 *in vivo*

Previous studies showed that GTP-bound human RALA and RALB bind the Exocyst subunits Sec5 and Exo84 in cultured cells and *in vitro* (Bodemann et al., 2011; Moskalenko et al., 2002; Moskalenko et al., 2003; Sugihara et al., 2002). To test whether *C. elegans* RAL-1 binds SEC- 5 and EXOC-8 *in vivo*, we used CRISPR-Cas9-dependent genome editing to insert sequences encoding fluorescent protein (FP) and epitope at the 3’ end of endogenous *sec-5* and *exoc-8* loci.

We then generated animals expressing SEC-5::mNG or EXOC-8::mNG together with our previously described mK2::3xFLAG::RAL-1 allele, without AID* (*re218*; Shin et al., 2018). As with mK2::RAL-1, SEC-5::mNG and EXOC-8::mNG were expressed ubiquitously and throughout the life cycle. These proteins were found predominantly in the cytoplasm but also showed plasma membrane enrichment, were they colocalized with RAL-1 (**Fig. 7; S4**).

**Figure 7:**
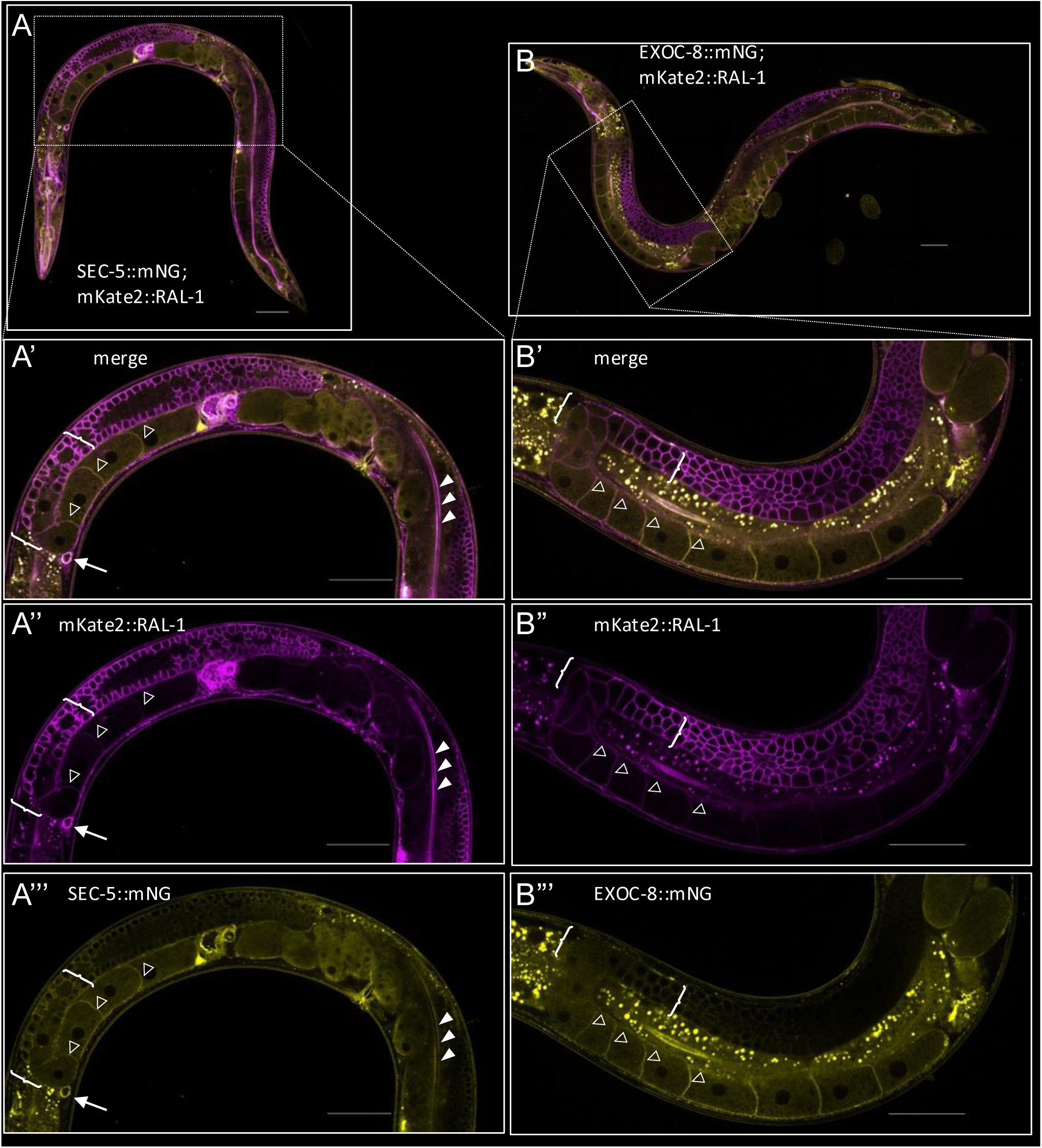
Endogenous RAL-1 colocalizes with endogenous SEC-5 and EXOC-8. A-A’’’) Whole animal **(A)** and zoomed images of merged **(A’)**, magenta **(A’’)** and yellow **(A’’’)** spinning disk confocal photomicrographs of endogenous mKate2::RAL-1 with endogenous SEC-5::mNG. Colocalization is indicated at coelomocyte plasma membrane (arrow), intestinal brush border/junctions (solid arrowheads), plasma membranes of neighboring oocytes (hollow arrowheads) and distal germline (brackets), among many other observed locales. **B-B’’’)** Whole animal **(B)** and zoomed images of merged **(B’)**, magenta **(B’’)** and yellow **(B’’’)** spinning disk confocal photomicrographs of mKate2::RAL-1 with EXOC-8::mNG. Colocalization is shown at intestinal lumen (solid arrowheads), plasma membranes of neighboring oocytes (hollow arrowheads) and distal germline (brackets), among many other observed locales. Scale bars = 50 µm.

### Co-immunoprecipitation Among RAL-1, SEC-5 and EXOC-8 Confirms Physical Association

Immunoblotting to detect epitope tags of endogenous mK2::3xFLAG::RAL-1, SEC- 5::mNG::2xHA, EXOC-8::mT2::2xMyx and EXOC-8::mNG::2xHA confirmed that tagged proteins were expressed at the expected sizes (**Fig. 8A-D**). To test whether these proteins physically associate *in vivo*, we performed co-immunoprecipitation (co-IP) assays in animals co-expressing tagged endogenous RAL-1 and tagged endogenous SEC-5 or EXOC-8. SEC-5::mNG::2xHA was immunoprecipitated with an α-HA antibody, resolved via SDS-PAGE, and EXOC- 8::mT2::2xMyc (mT2 = mTurquoise2, a blue FP) was detected in the pulldown using α-Myc antibody and repeated (**Fig. 8E,F**). Although the EXOC-8 signal was modest it was reproducible. Similarly, RAL-1 was detected in HA-based immunoprecipitations of both SEC- 5::mNG::2xHA (**Fig. 8G,H**) and EXOC-8::mNG::2xHA (**Fig. 8I,J**), respectively. These results indicate that RAL-1 associates with SEC-5 and EXOC-8 of the Exocyst complex *in vivo*.

**Figure 8:**
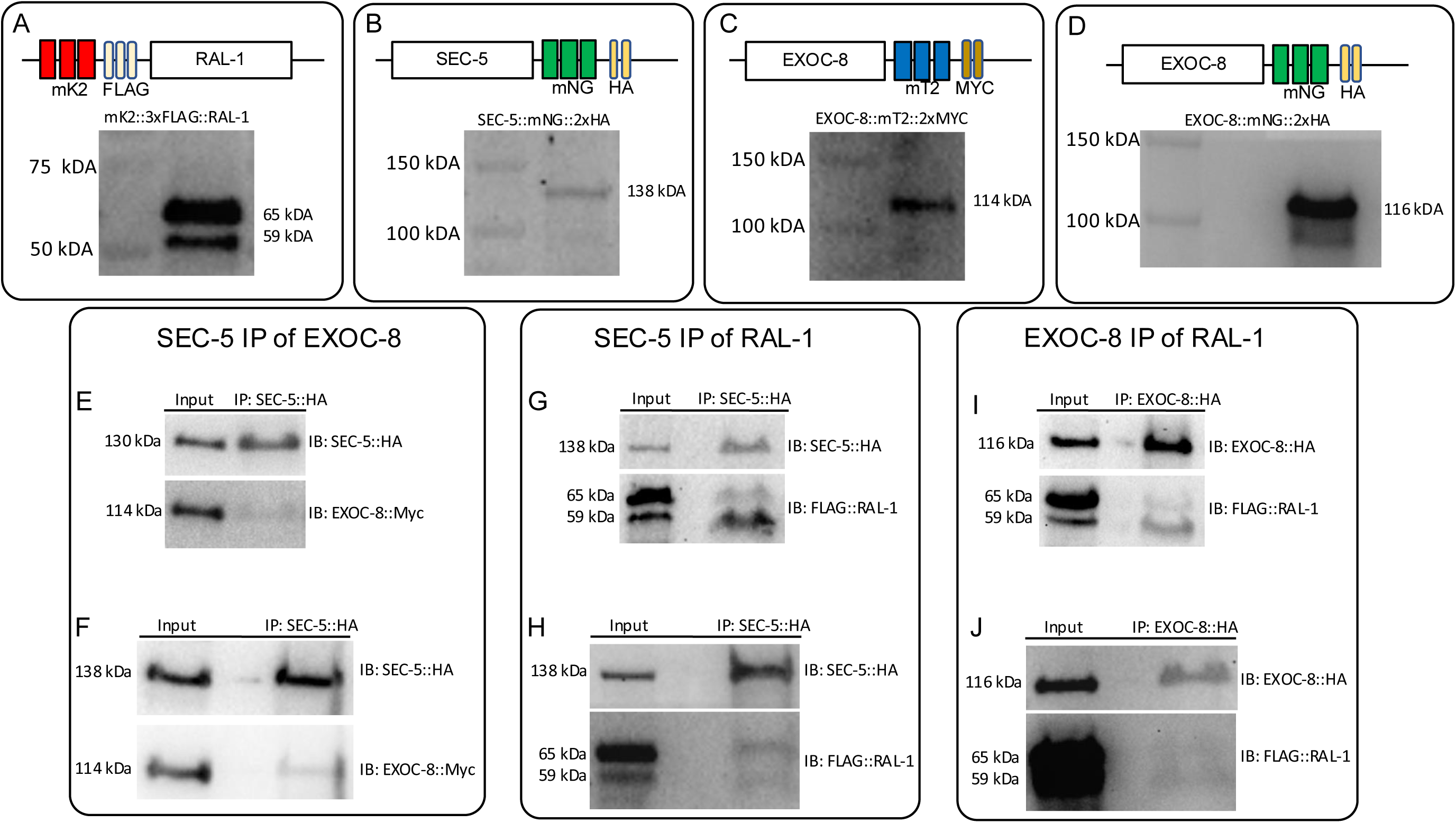
**Endogenous RAL-1 co-immunoprecipitates with endogenous SEC-5 and EXOC-8**. Detection via immunoblot and sizing validation of CRISPR-tagged endogenous **(A)** mKate2::2xHA::RAL-1 **(B)** SEC-5::mNG::2xHA, **(C)** EXOC- 8::mTurquoise2::2xMyc, and (**D)** EXOC-8::mNG::2xHA. **(E-J)** Immunoblots of immunoprecipitates and co-immunoprecipitates (Co-IPs) of endogenous proteins tagged with fluorescent protein and epitope. **(E)** Results of SEC-5::2xHA::mNG co- immunoprecipitating EXOC-8::2xMyc::mT2 and repeating **(F)**. **(G)** Results of SEC-5::2xHA::mNG co-immunoprecipitating 3xFLAG::mKate2::RAL-1 and repeating **(H)**. (**I**) Results of EXOC-8::2xHA::mNG co-immunoprecipitating 3xFLAG::mKate2::RAL-1 and repeating **(J)**.

### Structure-Guided Mutagenesis of RAL-1-binding residues in SEC-5 and EXOC-8

To test whether these interactions are direct and mediated by conserved binding interfaces, we introduced structure-guided point mutations into the endogenous *sec-5* and *exoc-8* loci. For SEC-5,we based our design on the crystal structure of human RALA with the Ral-binding domain of Sec5 (Fukai et al., 2003). Thr11 of Sec5, conserved in *C. elegans* and *Drosophila melanogaster* (**Fig. S5C**), was predicted to hydrogen bond to Tyr36 in RALA. Moreover, the T11A mutation dramatically decreased affinity of the Sec5 Ral-binding domain with RALA. Alphafold 2 modelling of *C. elegans* proteins predicted a similar interaction with SEC-5 Tyr16 and RAL-1 Tyr39 (**Fig. S5A-A’’’’**). Introducing the T16A mutation into SEC-5::mNG::2xHA via CRISPR-Cas9-dependent genome editing did not alter protein stability (**Fig. S5D**).

For EXOC-8, we similarly based our design on the crystal structure of human RALA with the putative Ral-binding domain of rat Exo84 (Jin et al., 2005). Ala228 and Lys233 of rat Exo84 are conserved in *C. elegans* and *Drosophila melanogaster* (**Fig. S5C**) and were both predicted to support binding to GTP-bound RALA via complex interactions but sterically inhibit binding with GDP-bound RALA. The space-occupying A228W and K233W mutations each dramatically decreased the affinity of human Exo84 Ral-binding domain with RALA (Jin et al., 2005). Alphafold 2 modeling of *C. elegans* proteins (**Fig. S5B-B’’’’**) predicted that A198W and K203W mutations in EXOC-8 would weaken associations with RAL-1, increasing distance between some interaction residues and abolishing interactions between others (**Fig. S5B,F**). Introducing the A198W mutation alone into EXOC-8::mT2::2xMyc via CRISPR-Cas9-dependent genome editing did not alter protein stability but had no biological effect, so was not studied further. Instead, we introduced the A198W,K203W double mutation into EXOC-8::mT2::2xMyc. The double mutation did not alter the stability of EXOC-8 (**Fig. S5E**) so we used the double mutation for further analysis.

### RAL-1 Binding-Deficient Mutants Show Synthetic Phenotypes

Neither the *sec-5(T16A)* nor *exoc-8(A198W,K203W)* single mutants displayed overt phenotypes. However, when combined with reduction-of-Exocyst function alleles in other Exocyst subunits, synthetic phenotypes emerged. *exoc-8(ok2523)*; *sec-5(T16A)* double mutants had significantly reduced body stature compared to single mutants (**Fig. 9A,B**). Similarly, *exoc-8(A198W,K203W); sec-5(pk2357)* M+Z- animals exhibited decreased stature (**Fig. 9C,D**), though all double mutants survived to adulthood. (Recall that the *sec-5(pk2357)* M+Z- single mutant survives to adulthood; **Fig. 2C**). *exoc-8(A198W,K203W)*; *sec-5(T16A)* double mutants also displayed reduced body size relative to single mutants and the wild type (**Fig. 9E,F**). These synthetic interactions suggest that the structure-guided mutations impair – but do not abolish – physical association between RAL-1 and the Exocyst complex *in vivo*.

**Figure 9:**
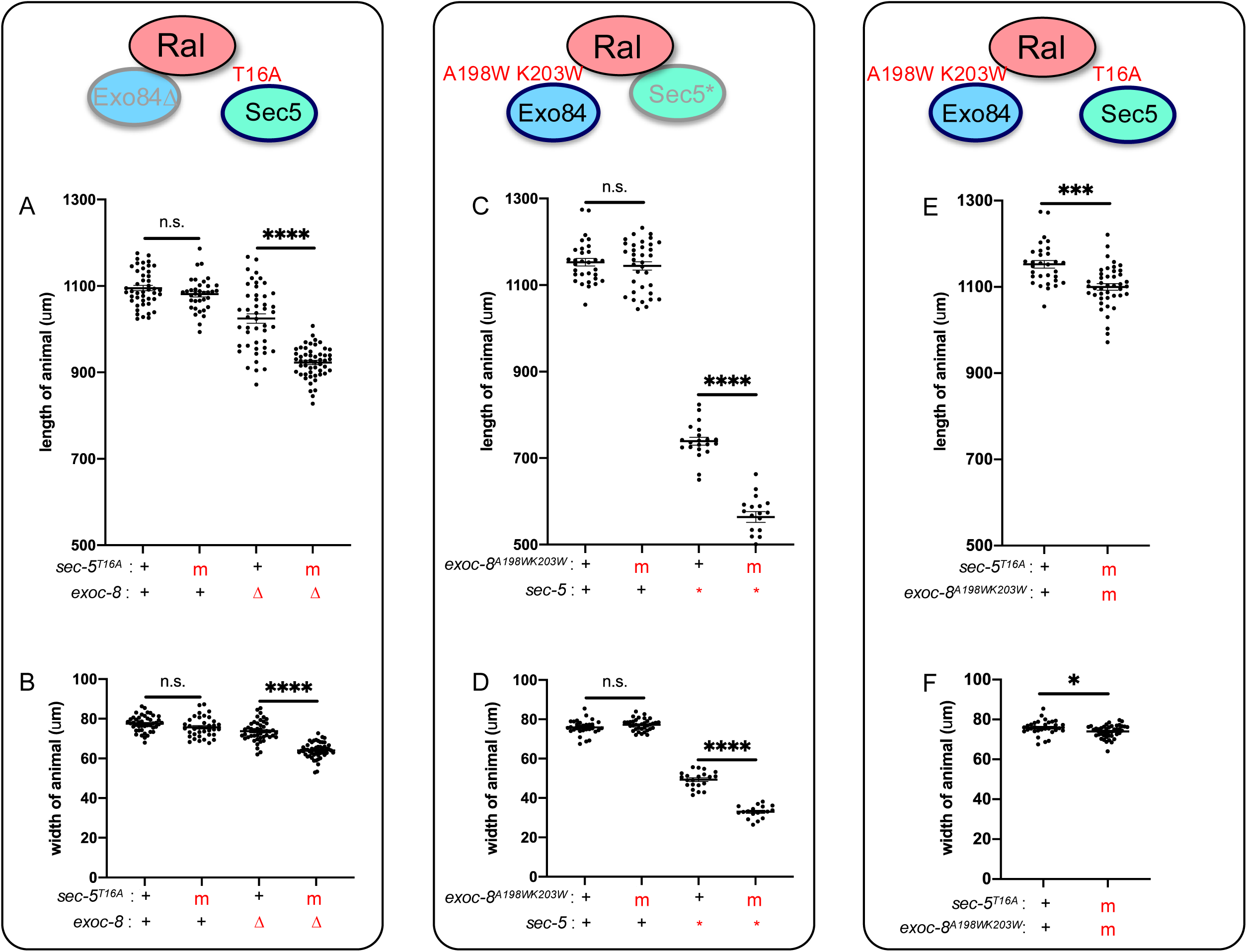
Structure-based putative Ral-effector interface mutations in SEC-5 and EXOC-8 reduce animal stature. A,B) The A198W,K203W double mutation in endogenous EXOC-8 does not alter animal stature in a wild-type background but aggravates both length and width defects of L4 M+Z- *sec-5(pk2357*\*) animals. **C,D)** The T16A mutation in endogenous SEC-5 does not alter animal stature in a wild-type background but aggravates both length and width defects of L4 M+Z- *exoc- 8(ok2523*Δ) animals. **E,F)** The double mutant between EXOC-8 A198W,K203W and SEC-5 T16A mutations confers decreased length and width. m = mutant, Δ = deletion, * = nonsense. *<0.05, ****<0.001, ****<0.0001, n.s. = not significant via *t* test.

Taken together, these results support the conclusion that RAL-1 physically and functionally associates with Exocyst subunits SEC-5 and EXOC-8 in *C. elegans*. These interactions are mediated by evolutionarily conserved binding surfaces and are functionally relevant for normal Exocyst activity *in vivo*. Combined with previous sections and mammalian studies, this establishes RAL-1 as a *bona fide* regulatory component of the Exocyst complex in metazoans.

## DISCUSSION

Our findings underscore the importance of studying small GTPases in intact systems. We developed an *in vivo* animal model to investigate the regulatory role of RAL-1, a Ras family small GTPase, in Exocyst-dependent vesicle tethering and exocytosis. While Ral GTPases have long been proposed to regulate the Exocyst complex, based on studies in mammalian cells and *in vitro* analysis, direct evidence for their physiological relevance in intact organisms has remained limited. Using only endogenous genes and proteins, we combined genetic, cell biological, phenotypic, and biochemical approaches to show that RAL-1 physically and functionally interacts with the Exocyst complex in *C. elegans*. RAL-1 is essential throughout post-embryonic development, genetically interacts with core Exocyst components, and is required for proper vesicle trafficking and dendritic arborization in PVD neurons. We find that RAL-1 acts both cell-autonomously in neurons and non-autonomously in the underlying epithelial cells that support dendrite arborization. Biochemically, RAL-1 binds directly to the Exocyst subunits SEC-5 and EXOC-8 through evolutionarily conserved surfaces, and structure-guided genome editing of these interfaces reveals synthetic phenotypes that highlight their biological importance. Together, these data establish *C. elegans* RAL-1 as a *bona fide* metazoan regulator of the Exocyst complex *in vivo* and *C. elegans* as a tractable animal model for dissecting Ral-Exocyst interactions at molecular, cellular, and developmental levels.

### How small GTPases regulate Exocyst function

The Exocyst functions as a conserved vesicle-tethering complex required for exocytosis in diverse systems. In the yeast *Saccharomyces cerevisiae*, where the Exocyst is well studied, its regulation is mediated primarily by the Rab-family GTPase Sec4p, along with additional inputs from the Rab GTPase Ypt31/32p in ER to Golgi trafficking, plus Rho families members Rho1/3p and Cdc42p for polarized secretion and vesicle trafficking via the actin cytoskeleton, and phospholipids PI(4)P and PI(4,5)P2 for membrane targeting (**Fig. S1A**; Volpiana et al., 2024; Wu et al., 2008). These small GTPases regulate specific Exocyst subunits to coordinate vesicle targeting, polarity, and membrane fusion. Notably, no Ral ortholog exists in fungi, and thus Exocyst regulation in yeast proceeds without branched Ral>Sec5 or Ral>Exo84 inputs into Subcomplexes 1 and 2, respectively. With this study we are confident that in metazoans, however, the Ral GTPases directly bind and regulate the Exocyst through interactions with the subunits Sec5 and Exo84. This evolutionary shift suggests that Ral-Exocyst interactions represent a metazoan-specific regulatory innovation, potentially allowing for more complex spatial and temporal control of exocytosis in multicellular context, perhaps through novel inputs.

Whether Ral functionally substitutes for specific Rab-family roles seen in yeast, such as the Sec4p role in binding and recruiting the Sec15p Exocyst subunit, is unclear. In *C. elegans*, the Rab proteins RAB-8 and RAB-10 have been implicated in polarized trafficking, both in the context of recycling endosomes and Exocyst function, raising the possibility of combinatorial or parallel regulation (Taylor et al., 2015; Zou et al., 2015).

Yet the general view of Rab GTPases in trafficking vesicles *vs.* Ras GTPases in signal transduction, are quite different models (Wennerberg et al., 2005; Reiner and Lundquist, 2018). Rab cascades function as ratchets that irreversibly advance organelles through defined states (Barr, 2013). In contrast, Ras signaling is better understood as a dose-dependent regulator, like a rheostat. Different strengths and durations of Ras activation are decoded into distinct biological outcomes, providing a graded, analog mode of regulation rather than a directional switch (Bustelo et al., 2018; Li et al., 2018). Ral is a Ras family GTPase: does it intersect differently with vesicle trafficking than do Rab proteins? Or does the same protein regulate vesicle trafficking differently than it does with signal transduction? This distinction raises the intriguing possibility that Ral GTPases integrate signaling pathways with membrane trafficking using mechanisms distinct from Rab-mediated control. To speculate, perhaps the difference is imposed by the distinct GDIs (Guanine Dissociation Inhibitors, intercompartmental trafficking proteins for small GTPases) used by Rab vs. Ras proteins. Interestingly, a no GDI that functions for Ral has been reported.

Our results underscore this emerging view. RAL-1 interacts physically and functionally with the Exocyst complex and appears to be required for multiple steps in Exocyst-dependent trafficking, including vesicle formation or cargo loading, transport, tethering, and likely fusion. The requirement for RAL-1 both within dendrites and in adjacent epithelium further suggests a broader regulatory influence, possibly coordinating polarized trafficking across multiple cell types. As such, RAL-1 may represent a metazoan-specific mode of Exocyst regulation, distinct from Rab- and Rho-based mechanisms observed in unicellular eukaryotes.

### Requirements for Ral>Exocyst in a live animal

At first glance, one might assume that the Exocyst complex is essential for cell viability, given the fundamental requirement for trafficking of integral membrane proteins and secreted factors in multicellular organisms. This expectation is initially supported by findings in cultured cells, where the Ral>Exocyst pathway is essential, and in animals models: M+Z- mutants of *Drosophila* Ral die during embryogenesis or early larval stages, and double knockouts of RalA and RalB in mice result in early embryonic lethality (Balakireva et al., 2006; Peschard et al., 2012; Wang et al., 2015). Yet remarkably, in investigation of the mouse RALA and RALB single and double knockouts the word “Exocyst” is not mentioned once (Peschard et al., 2012). In *C. elegans*, knockout of *ral-1* in early embryonic cells causes a failure to recruit the Exocyst to the plasma membrane (Armenti et al., 2014a). Consistent with these findings, our studies reveal that partial loss of Exocyst function disrupts diverse developmental events, including severe defects in dendritic arborization, failure of germline development, and morphogenetic abnormalities such as vulval rupture in M+Z- *sec-5* mutants.

Given these observations, it is surprising that complete depletion of RAL-1 or core Exocyst components via the ZF1/ZIF-1 degron system results in only partial embryonic lethality, with many animals surviving to L1 larval stage (Armenti et al., 2014a), though it is striking that single-cell complete depletion of RAL-1 in polarized secretory cells causes defects as extreme as similar depletion of CDC-42 (Abrams and Nance, 2021). This observation suggests that embryos can progress through development farther than expected in the absence of Ral>Exocyst activity, raising the question of how vesicle trafficking is sustained.

One plausible explanation is partial redundancy between the Exocyst-dependent direct secretion and other secretory pathways, particularly recycling endosomes. This idea is supported by prior observations in *C. elegans*, where RAL-1 and Exocyst components appear genetically redundant with RAP-1/Rap1, another Ras family small GTPase, for embryonic morphogenesis. In embryos, simultaneous deletion of RAP-1 and RNAi-dependent depletion of RAL-1 and Exocyst components leads to disorganized adherens junctions, defective localization of HMP-1/⍺-catenin to the basolateral plasma membrane, and dead embyos (Frische et al., 2007). More recently, RAP-1 and its activating GEF, PXF-1, were implicated in the fission and release of recycling endosomes (Rodriguez-Polanco et al., 2023), a role that could functionally parallel or compensate for Exocyst-mediated tethering. Together, these findings suggest that RAL-1/Exocyst and RAP-1/recycling endosome pathways may define two partially redundant routes for vesicle targeting and secretion. Loss of both pathways may be incompatible with development, explaining why embryos lacking both RAL-1 and RAP-1 die earlier than those lacking either alone. And thus the Exocyst may not be required for secretion via recycling endosomes.

In conclusion, our findings establish RAL-1 as a core regulator of Exocyst-mediated vesicle trafficking in a live, developing animal. Through precise genetic manipulation and tissue-specific depletion, we demonstrate that RAL-1 functions both cell-autonomously within neurons and non-autonomously in surrounding epithelium to direct vesicle delivery essential for dendritic patterning. These interactions are mediated by evolutionarily conserved interfaces between RAL-1 and key Exocyst subunits, and their disruption produces synthetic phenotypes that underscore their physiological significance. More broadly, our work reveals that the Exocyst is not simply a terminal tethering machine but integrates with small GTPase signaling to coordinate cargo assembly, transport, and secretion. The unexpected developmental resilience to loss of RAL-1 or Exocyst activity further suggests compensatory roles for other trafficking systems, such as recycling endosomes, highlighting a complex, layered architecture of membrane traffic in metazoan cell biology. Together, these results provide *in vivo* validation of a Ral>Exocyst signaling axis, define key principles by which small GTPases regulate polarized secretion in intact tissues, and establish a genetically and developmentally tractable *in vivo* system for investigating Exocyst function

## METHODS

### *C. elegans* handling and genetics

Strains used in this study were derived from the N2 Bristol wild type. Animals were grown at 20°C on NGM plates spotted with OP50 *E. coli* unless otherwise noted. Strains used in this publication are listed in Table S1, Oligonucleotides in Table S2, and Plasmids in Table S3.

### CRISPR/Cas9-dependent genome editing

We selected guide RNAs that satisfy three approaches. First, G and not T nucleotides were selected at positions -1/-2, or GCGG and not T-TT nucleotides at positions -1/-2/-3/-4 (Farboud and Meyer, 2015; Wang et al., 2014). Second, we considered strong predicted specificity and efficiency scores using the CRISPOR (http://crispor.tefor.net/) algorithm, which incorporates the original MIT specificity score. Third, we considered strong predicted efficiency scores using the WU-CRISPR (http://crisprdb.org/wu-crispr/) algorithm.

CRISPR-Cas9 injection mixes were prepared essentially as described (Fakieh et al., 2022; Ghanta et al., 2021). All preparation steps were performed on a dedicated RNase-free bench. Final concentrations were calculated per 20 μl reaction. To assemble the mix, 1 μl of 5 μg/μl *Streptococcus pyogenes* Cas9 protein (PNAbio, #CP01) was added to yield 0.25 μg/μl final concentration. This was combined with 1 μl of 2 μg/μl stock tracrRNA (IDT) for a final concentration of 0.1 μg/μl. Gene-specific crRNAs and *dpy-10* co-CRISPR crRNA were added as 1.4 μl each of 0.4 μg/μl stocks, yielding 0.028 μg/μl final concentration per crRNA. The pre-final mix was incubated at 37°C for 15 minutes to anneal crRNAs to tracrRNA and assemble the ribonucleoprotein complex. Following incubation, 3.3 μl of 20 μM stock *dpy-10(cn64*gf) ssODN repair oligo was added for a final concentration of 3.3 μM. Gene-specific repair templates were generated by PCR, purified, and subjected to denaturation/renaturation as described (Ghanta and Mello, 2020), then added at ≤300 ng/μl stock for a final concentration of 100 ng/μl, ensuring the total PCR concentration did not exceed 2 μg/20 μl to avoid viscosity issues. Finally, nuclease-free water was added to bring the total volume to 20 μl.

Adult animals were selected for microinjection based on carrying no more than a single row of embryos in the uterus. Injections were carried out as described previously (Kadandale et al., 2009). Injected P0 animals were transferred singly to spotted plates and incubated at 20 °C for ∼84-96 hrs. F1 progeny expressing the *dpy-10(cn64*gf) Rol phenotype (Arribere et al., 2014) were identified and transferred either singly or in pairs to plates. Once F1 Rols had produced F2 embryos, the parental F1 animals were picked to PCR tubes, lysed, and analyzed by single-worm PCR. Homozygous F2 progeny were obtained by selecting non-Rol siblings segregating from confirmed knock-in F1 parents. All edited loci were validated by Sanger sequencing across the region of homology-directed repair (HDR), and only animals carrying the precise, expected edits were retained for further study. Aberrant or unintended mutant alleles were discarded.

### Imaging

Live animals were mounted on 3% agar pads in 5 µl of 2 mg/ml tetramisole in M9 buffer. DIC and fluorescent micrographs were captured using a Ti2-Nikon inverted microscope equipped with a Yokogawa CSU-W1 Spinning Disk and a Photometrics Prime BSI camera.

Green fluorescence (mNG) was captured 488 nm, red fluorescence (mK2) was captured at 561 nm, and blue fluorescence (mTagBFP2) was captured at 405 nm. NIS Elements Version 4.30 software was used to capture and process images.

### Quantifying animal defects

Synchronized embryos were obtained by picking adults (24 hours post-late L4) onto plates with food and picking off 1 hour later. Developmental endpoints of growth defects were determined among resulting animals: embryos 24 hours later, larvae 48 hours later and adults 72 hours later. Vulval rupture was scored by picking L4 animals to plates and scoring 24 hours later.

Animal size was quantified using synchronized adults 72 hours after egg laying. Animals harboring sec-5(*) were analyzed at 48 hours after egg laying, as mutant adults ruptured. Synchronized animals were mounted on 3% agar pads on slides. DIC photos were taken using a spinning disk confocal microscope with a 40x objective lens. Length and width of animal was analyzed using NIS Elements Version 4.30 software. Length was measured as distance from the anterior to posterior (A-P) end and width as the ventral to dorsal (V-D) at the A-P midpoint where the vulva is located.

For quantifying defects in PVD dendritic arborization, mid- to late-L4 animals harboring the *wdIs52[F49H12.4p>GFP + unc-119(+)]* transgene expressing GFP in PVDs were mounted in M9 buffer on 3% agar pads with anesthetic. Using the 40x objective on the spinning disk confocal microscope, we captured z-stack images of PVD dendrites for the side – left or right – facing the objective. Anterior and posterior sections of the PVD were defined as before and after the cell body. We counted numbers of secondary, tertiary and quaternary dendrite branches from the resulting collapsed z-stacks. The DMA-1::GFP exocytic cargo marker was scored in mid- to late- L4 animals mounted and imaged as above. Number of punctae in primary dendrites was counted in the segment 50 µm to 100 µm from the cell body. Intensity of punctae was measured 50 µm from the cell body using NIS Elements Version 4.30.

### Depletion of gene product

RNAi was performed as previously described (Shin et al., 2018). Briefly, 20 µl of HT115 E. coli was seeded on NGM plates with carbenicillin and tetracycline. L4 animals were picked to plates the following day, transferred 24 hours later, and removed after an additional 24 hours. *Luciferase(RNAi)* (Shin et al., 2018) and *pop-1(RNAi)* were used as negative and positive controls, respectively. Phenotypes were only scored on those experiments with 100% lethality with *pop-1(RNAi)*. RNAi experiments were performed in 23°C.

Auxin-inducible degron (AID*; Zhang et al., 2015) sequences (135 bp) were inserted into the N-terminal mKate2::3xFLAG tag of RAL-1 (Fakieh et al., 2022). The resulting AID::mKate2::3XFLAG::RAL-1 was crossed into strains harboring transgenes expressing the TIR1 cofactor ubiquitously or behind neuronal-specific or hypodermal-specific promoters (Ashley et al., 2021). Plates with synchronized animals were treated with 500 µl 1 mM auxin (indole-3- acetic acid; IAA; diluted from 400 mM auxin stock in ethanol, as described (Fakieh and Reiner, 2025). Vehicle control animals were treated with same concentration of EtOH.

### Protein detection

For immunoblotting, mixed stage animals were washed from plates using M9 buffer and lysed in 4% SDS loading buffer by boiling at 90°C for 5 minutes. Protein samples were run on 4%–15% SDS gel (Bio-Rad) and blotted on Immobilon-P Membrane, PVDF (EMD Millipore, IPVH00010). Anti-FLAG antibody (Sigma-Aldrich F1804), anti-HA antibody (Proteintech 51064-2-AP), anti-MYC antibody (Cell Signaling Technology 9B11) and anti-⍺- tubulin antibody (Sigma-Aldrich T6199) were diluted 1:2000 in blocking buffer (6% w/v non-fat dry milk in PBST). HRP-conjugated goat anti-mouse secondary antibody (MilliporeSigma 12- 349) was diluted in 1:5000 in blocking buffer. Chemiluminescent detection was performed using ECL reaction (Thermo Fisher Scientific) and detected using the Bio-Rad Chemidoc MP Imaging System Hood III and film processor, SRX-101A (Konica Minolta) on X-ray film (Phenix). For co-immunoprecipitation, animals as above were lysed using a sonicator (QSONICA Q500) and incubated with anti-HA antibody (Proteintech 51064-2-AP) and Dynabeads protein A (Invitrogen™ #10002D) washed by PBST and 0.1% 1M glycine pH=1 in 4°C overnight and detected by western blot as described.

### Software and statistical/data analysis

Published crystal structures were examined using Pymol (version 2.5.4). PDB accession numbers: 1U90, 2KE5, 5YFP and 6P0J. Protein models of RAL-1 interaction with SEC-5 and EXOC-8 were predicted by Alphafold2 through ColabFold v1.5.5 (Mirdita et al., 2022) and displayed by Pymol (version 2.5.4). Molecular distances between residues in RAL-1 and EXOC-8 were measured by UCSF ChimeraX (version 1.7.1). Data analysis and graphing was performed using Prism 9 (version 9.0.1) and Microsoft Excel (version 16.45).

## ACKNOWLEDGEMENTS

We thank L. Vergara in the Center for Advanced Imaging for technical assistance, Zheng Zhou, Buck Samuel , and Michael Galko for helpful discussions, N Rasmussen, RP Banerjee and T Duong in the Reiner lab for technical expertise and members of the Reiner lab for helpful discussions. We thank Andrew Golden (NIDDK) for sharing his composite CRISPR protocol, David Sherwood and Monica Driscoll for sharing strains. Some strains were provided by NBRP, which is funded by the Japanese government. Some strains were provided by the *Caenorhabditis* Genetics Center, which is funded by the NIH Office of Research Infrastructure Programs (P40 OD010440). Wormbase was used routinely (Sternberg et al., 2024). This work was funded by grants R35 GM144237 and R03 CA289854 to D.J.R.

---

**Figure S1:**
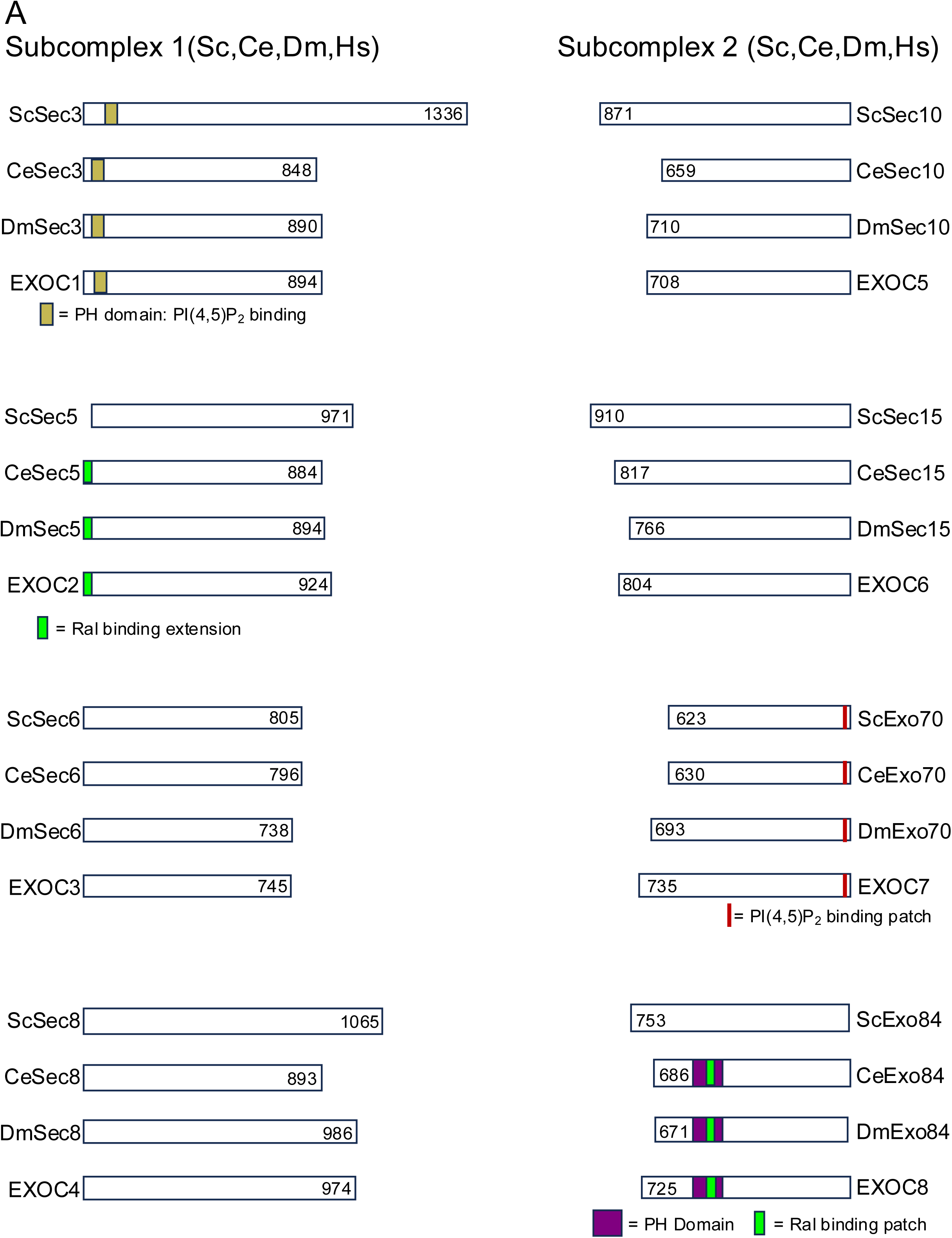
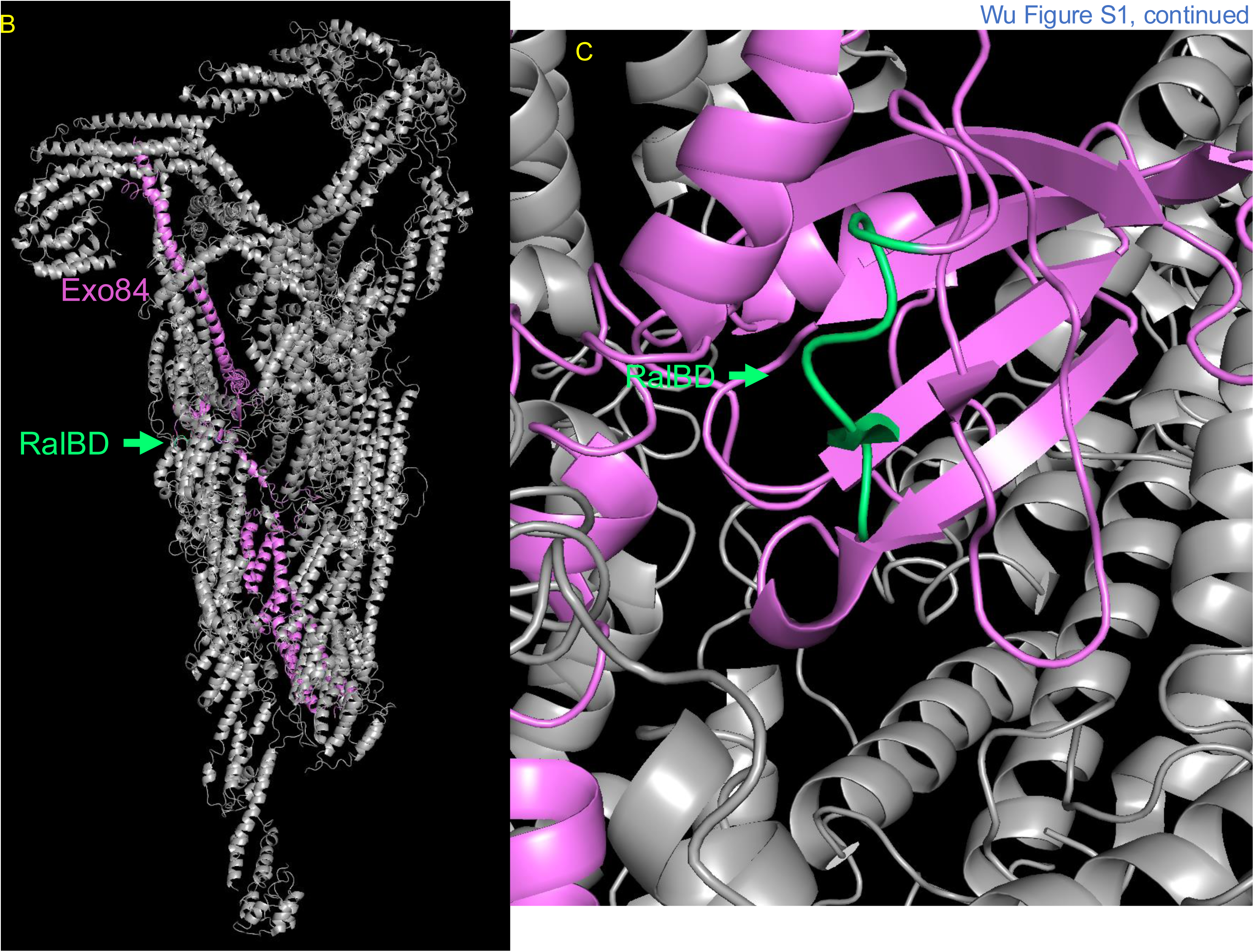
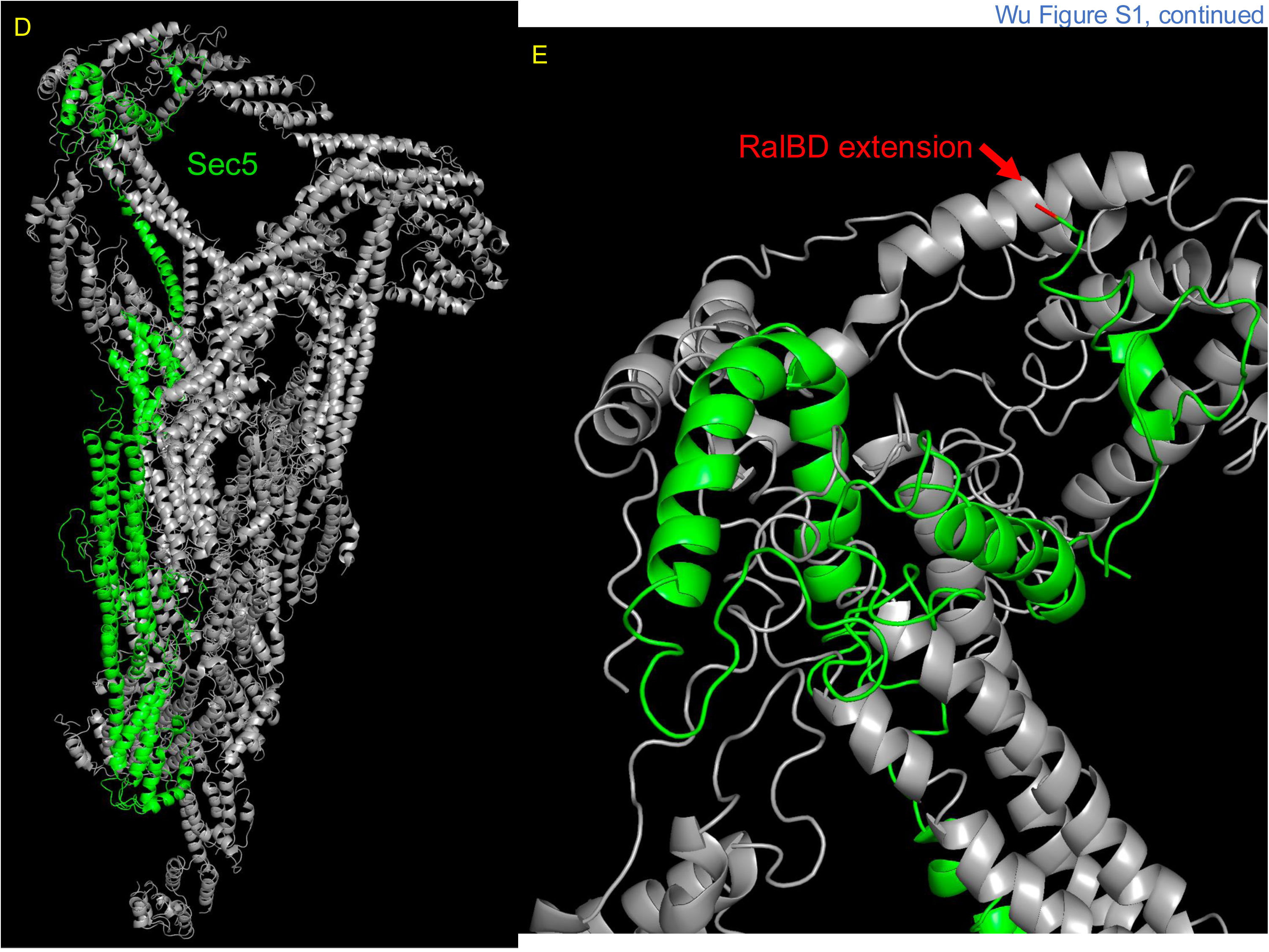
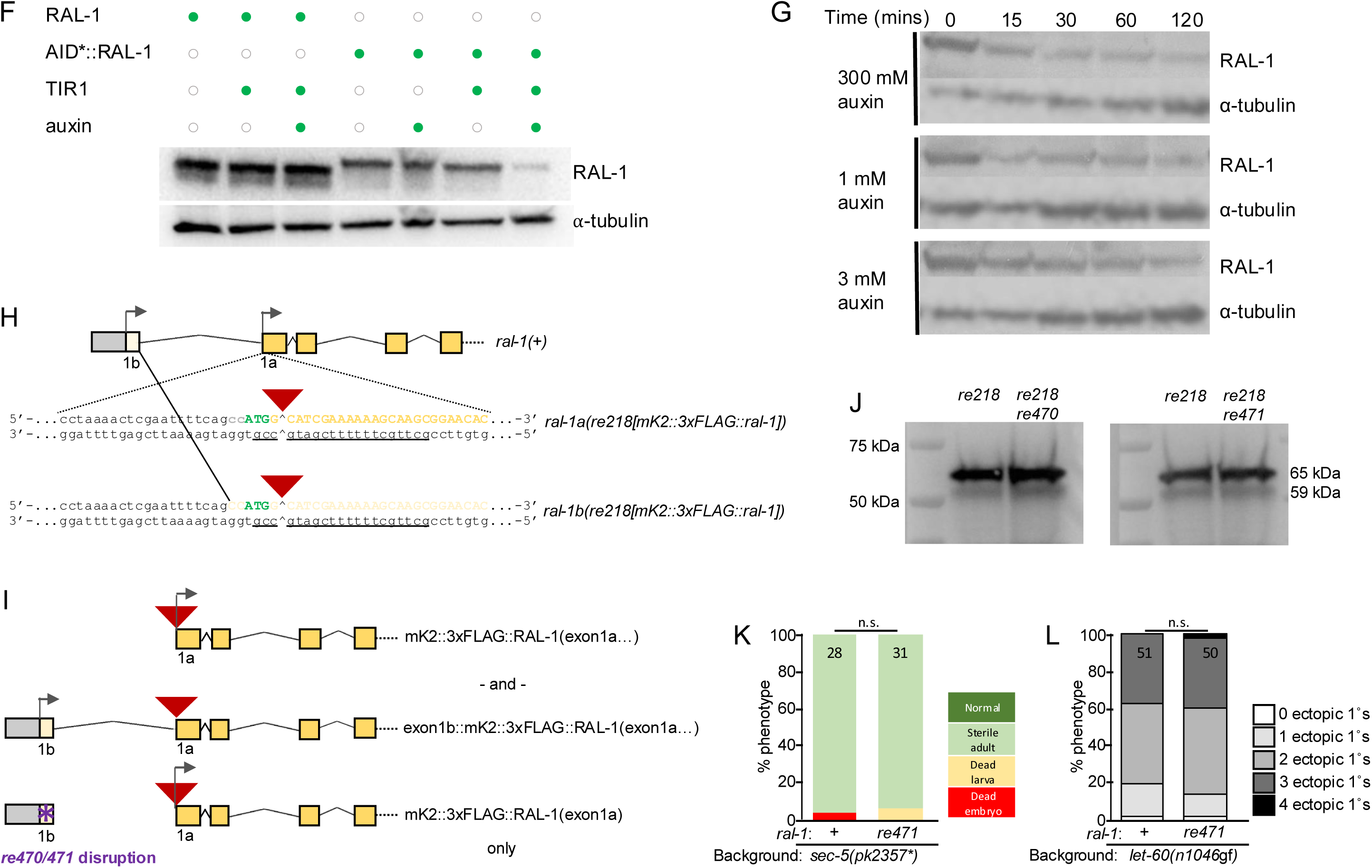
Ral, the Exocyst complex, and conserved sequence domains and motifs in the eight Exocyst subunits from four species. **(A)** *S. cerevisiae* (Sc), *C. elegans* (Ce), *D. melanogaster* (Dm), and *H. sapiens* (Hs), divided into Subcomplex 1 and Subcomplex 2. The Exocyst is a CATCHR (<u>c</u>omplexes <u>a</u>ssociated with tethering <u>c</u>ontaining <u>h</u>elical <u>r</u>ods) family of multisubunit tethering complexes (MTCs) used in vesicular trafficking and fusion. Exocyst subunits do not contain catalytic domains. Rather, all of them are composed primarily of alpha helices with limited regions of binding interfaces. Many of these alpha helices are organized into helical bundles. Each subunit in turn is assembled into pairs with its partner (Sec3-Sec5, Sec6-Sec8, Sec10- Sec15, Exo70-Exo84), the partners are assembled into Subcomplexes 1 and 2 (Sec3/5/6/8 and Sec10/15/Exo70/84, respectively), and the two subcomplexes assembled into the heterooctameric mature complex (Mei *et al*. 2018). Consequently, the helical bundles in part govern complex assembly and are thus distinctive for each subunit: the are distinctive and the sequence and structural levels and can be identified among orthologs of a subunit across evolution. For example, the Pfam Sec3 domain is poorly conserved at the level of primary sequence but is readily identified through sequence analysis programs like SMART, used here. The eight Pfam domains of helical clusters, one for each subunit, are not shown in this figure, but are shown in Appendix 1 for this study. This Appendix provides all the sequence files and alignments where domains of each protein are annotated and alignments of each subunit family are collated. Outside of helical bundles, the conventional domains of the Exocyst are shown in this schematic: an unconventional PH domain in Sec3 validated in yeast as binding the phospholipid PI(4,5)P, and an unconventional PH domain in Exo84 that no longer binds lipids (and is absent in yeast). Studies of Ral have implicated two binding sites, in Exo84 and Sec5.These were originally identified via yeast two hybrid and validated with fragment binding *in vitro* and in cultured cels. The fragments of mammalian subunits have been crystalized with human RALA (Fukai 2003; Jin 2005). The Sec5 fragment is an N-terminal extension present in metazoans but not in yeast. The Exo84 fragment includes a non-functional mid-protein PH domain, potentially evolved to support Ral binding. The corresponding putative Ral-binding sequence in Exo84 is not conserved in yeast. Alignments of both are shown in **Fig. S8C**. Subcomplex 1 subunits: Sec3: Uniprot #s: P33332, Q20678, Q9VVG4, Q9NV70 Sec5: Uniprot #s: P89102, Q22706, Q9VQQ9, Q96KP1 Sec6: Uniprot #s: P32844, Q19262 , Q9V8K2, O60645. Sec8: Uniprot #s: P32855, Q9XWS2, Q9VNH6, Q96A65. Subcomplex 2 subunits: Sec10: Uniprot #s: Q06245, Q18406, Q9XTM1, O00471 Sec15: Uniprot #s: P22224, Q18286, Q9VDE6, Q8TAG9 Exo70: Uniprot #s: P19658, P91149, Q9VSJ8, Q9UPT5 Exo84: Uniprot #s: P38261, Q95Q35, Q9VBI4, Q8IYI6 Pfam domains for each Exocyst subunit: detectable by sequence-based algorithms, structural conservation, shared across orthologous proteins, thought to be a core complex module **(B)** A model of yeast Exocyst complex structure derived from Cryo-EM (Mei *et al*. 2018). Structure was analyzed using PyMOL 3.1.6.1. Exo84 is colored pink. The Pfam Exo84 domain, organized into helical bundles, is evident at the bottom of the model (see Appendix for annotated individual sequences for all four species for Exo84, at the C-term, bottom here, but also for other subunits. **(C)** A zoom to see the loop corresponding to the Ral-binding domain (RalBD) of Exo84, highlighted and indicated by an arrow in blue-green. This loop is present in yeast Exo84, of which this is a model, but the sequence is only that of the RalBD in metazoans (Appendix; **Fig. S1A Fig. S8C**). **(D)** The same model based on CryoEM model of the Exocyst, rotated 180° on its long axis, with Sec5 colored lime green. **(E)** A zoom to see the N-terminus of Sec5 at the top of the model, with the N-terminal methionine colored red for orientation. Metazoans encode an N-terminal extension at this point, not found in yeast, that is thought to contain another RalBD (Appendix; **Fig. S1A Fig. S8C**). **(F)** Immunoblot validation of chemical genetic depletion of AID*::mKate2::3xFLAG::RAL-1: animals of genotype *ral-1(re218[mKate2::3xFlag::ral-1])* or *ral- 1(re218re319[AID*::mKate2::3xFlag::ral-1])* singly or in combination with *ieSi57[eft-3p>TIR1::mRuby]*, with or without addition of auxin (IAA = indole acetic acid) and immunoblotted with ⍺-FLAG antibody. The upward band shift between the first three and the latter three lanes is due to the addition of AID*. **(G)** Immunoblot detection of time course of auxin addition and different concentrations indicates that depletion of RAL-1 is incomplete regardless of dose and time. This may be due to inaccessibility of the AID* tag to TIR1-auxin when bound in complexes, specifically the Exocyst, or perhaps poor accessibility of the AID* target in the context of tagged RAL-1. **(H)** Note the two RAL-1 bands variably visible in most exposures. In our previous study (Shin *et al*., 2018, Fig. S6), we hypothesized that the upper band is the predicted 61 kDa RAL-1B isoform with a non-canonical-length N-terminal extension unique to Nematoda, while the lower band is the 56 kDa RAL-1A isoform with a canonical-length N-terminal extension present in most metazoans. The alternative splicing of the upstream exon 1b and canonical exon 1a are shown: both are predicted to splice into the RAL-1 tags: AID*::mK2::3xFLAG or mK2::3xFLAG tag and thus be detectable by western blot. **(I)** We selectively disrupted exon 1b with CRISPR-based STOP-IN (Wang, 2018), generating alleles *re218re470* and *re218re471*. We evaluated the function of deficient exon 1b animals in the two known activities of *ral-1*: exocyst and signaling. **(J)** Neither *re470* nor *re471* altered expression of tagged endogenous RAL-1 as detected by ⍺-FLAG immunoblotting. Since by RNAseq RAL-1A is predicted to be much more abundant than RAL-1B (see Results), RAL-1A may mask RAL-1B protein as well as the consequences of disrupting exon 1b. Bands sizes remain mysterious. The 65 kDa band corresponds to unmodified RAL-1B or perhaps post-translationally modified RAL-1A. The much less abundant 59 kDa band corresponds to the expected size of unmodified RAL-1A. **(K)** *re471* did not alter Exocyst function in animals deficient for M+Z- *sec-5 (*this study) or **(L)** induction of 2° VPC fates as assayed by antagonism of *let-60(n1406*gf) induction of ectopic 1° VPCs (our other studies). We conclude that, in addition to be an apparently nematode-specific alternative exon of *ral-1*, exon 1b is not required for established functions of *ral-1* in Exocyst or signal transduction.

**Figure S2.**
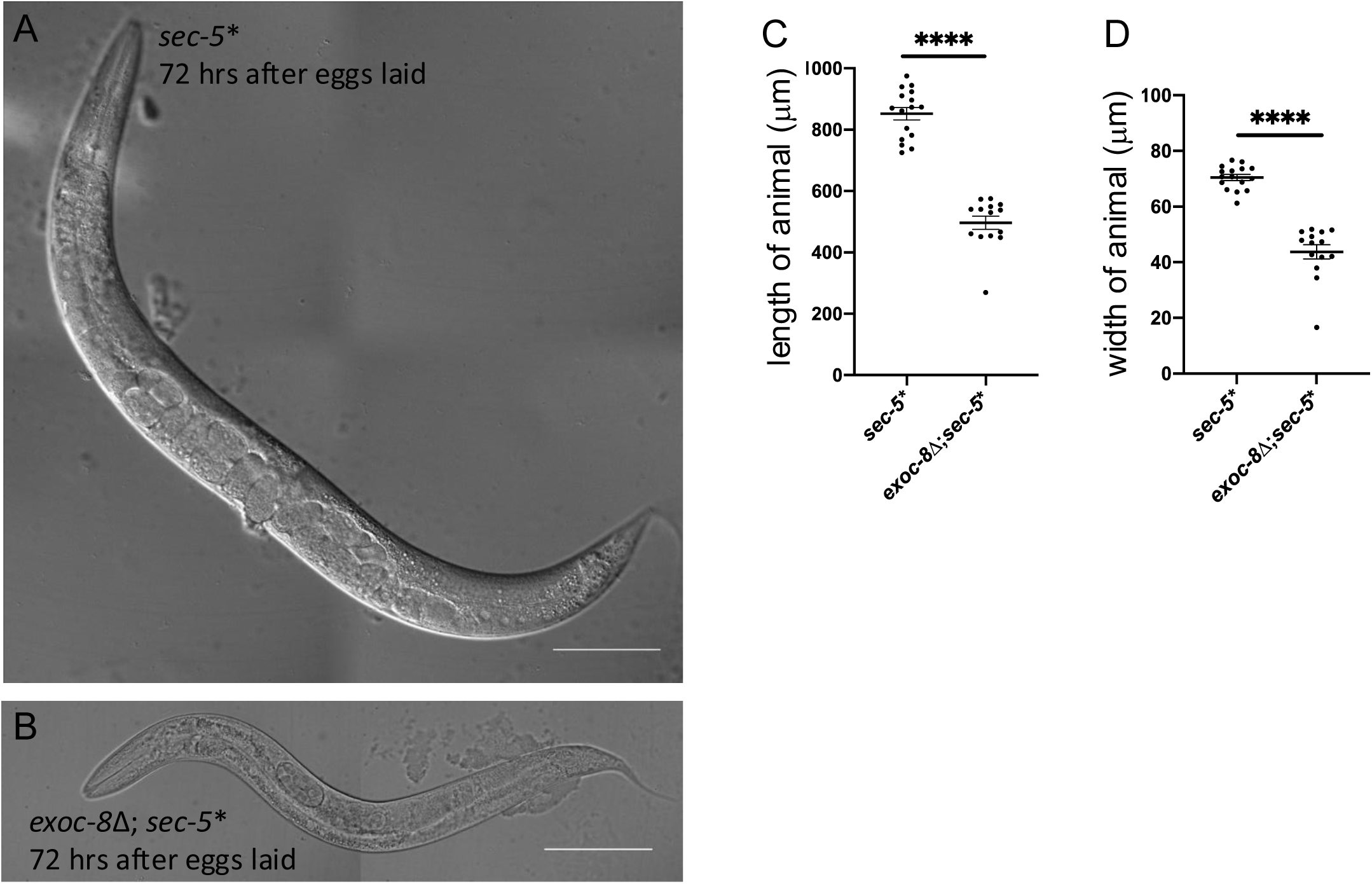
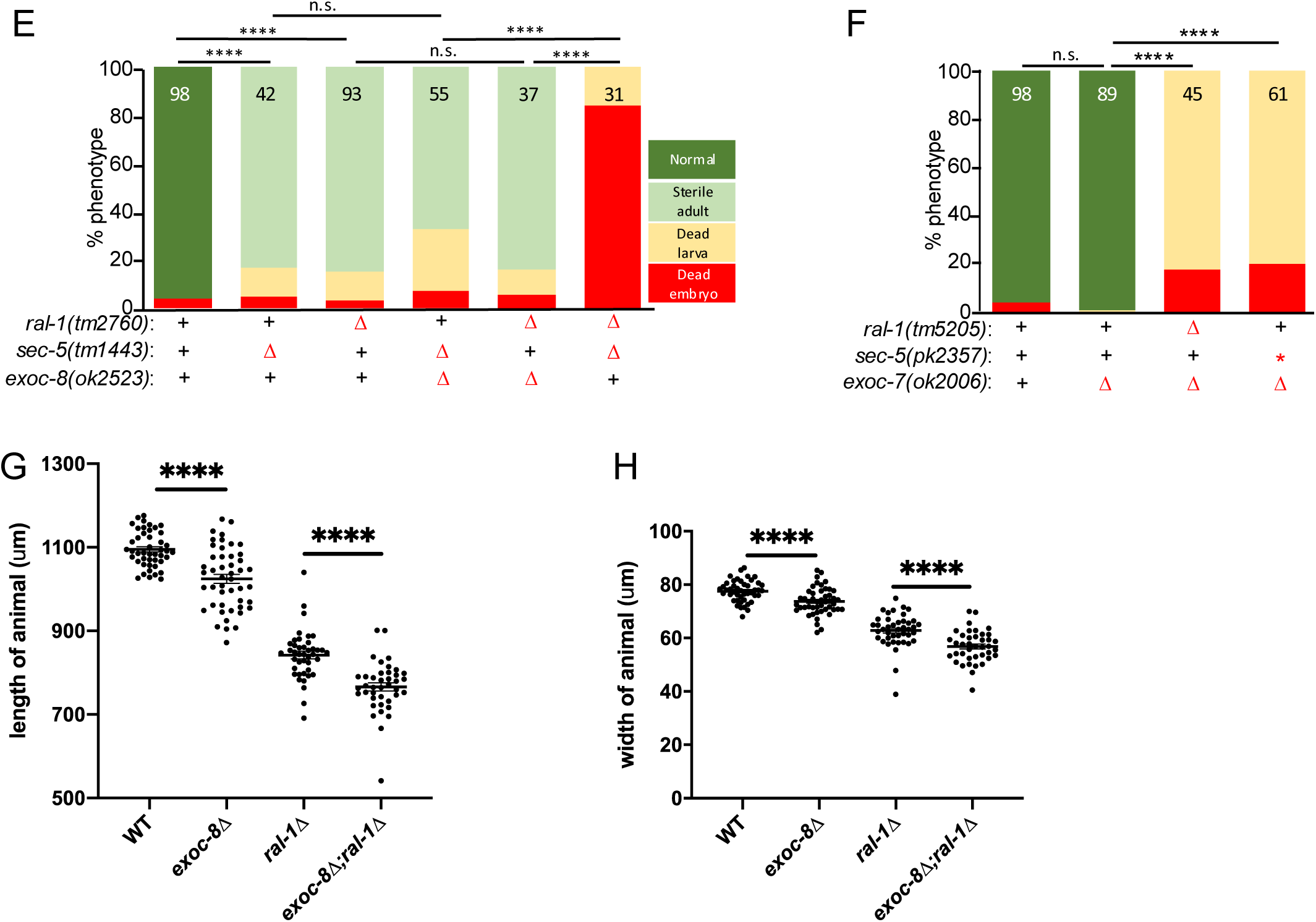
Combining reduction-of-function mutations among *ral-1* and Exocyst components aggravates growth/developmental effects. A-D) *exoc-8(ok2523)*; *sec-5*(*pk2357*\*) double mutants from Fig. 1C did not die at earlier larval stages but were substantially decreased in length **(C)** and width (**D)**, which could be superficially mistaken for earlier lethality. **(A,B)** Representative DIC photomicrographs of synchronized *sec-5*(*pk2357*\*) **(A)** and *exoc-8*(*ok2523)*; *sec-5*(*pk2357*) **(B)** animals. **(E,F)** Genetic interactions among second alleles of *ral-1*, *sec-5* and non-essential subunit-encoding *exoc-7* in support of Fig. 1C. Color-coded genotype key is right of the two graphs. **(E)** Genetic interactions with second alleles *ral-1(tm2706)* (Zand *et al*., 2011) and *sec- 5(tm1443)* with the *exoc-8(ok2523)* allele used in Fig. 1C. (F) *sec-5* and *ral-1* alleles used in Fig. 1C but in combination with deleted *exoc-7*, which encodes the Exo70 non-essential subunit reported to dimerize with Exo84 and assemble with Subcomplex II (Mei 2018; LePore 2018). (G,H) *exoc-8(ok2523)* and *ral-1*(*tm5205)* single and double mutants from Fig. 1C did not die at earlier larval stages but manifested substantially decreased length (**G**) and width (**H**), which could be superficially mistaken for earlier lethality.

**Figure S3.**
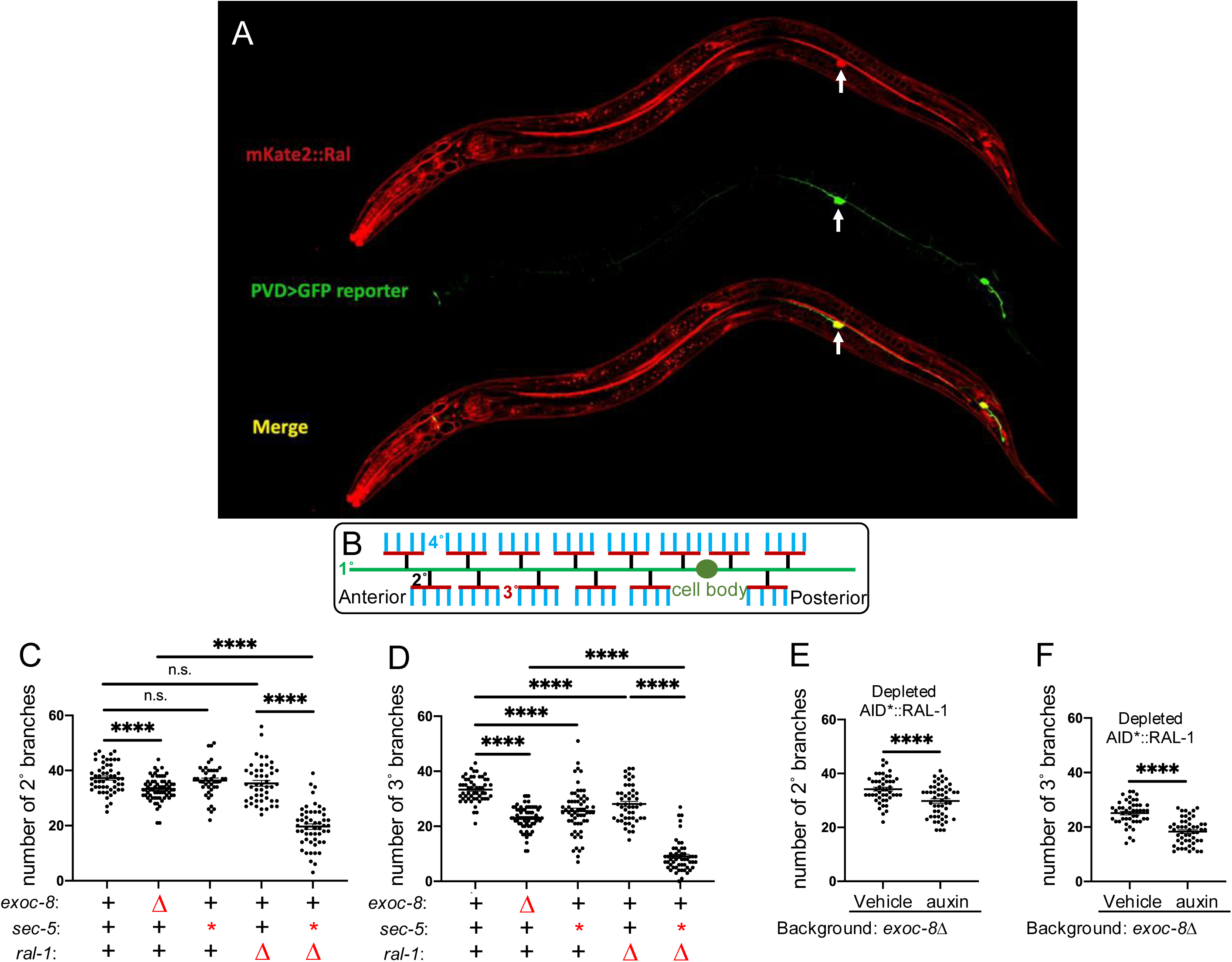
RAL-1 is required for full exocyst-dependent PVD dendritic arborization. **(A)** Maximum intensity projection of z-series confocal photomicrographs of endogenous mKate2::3xFLAG::RAL-1 protein colocalized with GFP-labelled (*wdIs52[ser-2(prom3)p>gfp])*) cell body of the PVD neuron (arrows); red = RAL-1, green = PVD label, yellow = merge. **(B)** A schematic of the 1° (green), 2° (black), 3° (red) and 4° (blue) branches of the PVD neurons. Anterior is left for both animals. **(C,D)** The *exoc-8(ok2523)*; *ral-1(tm5205)* double mutants exhibit more severe defects in 2° **(C)** and 3° **(D)** dendritic branching than each single mutant. **(E,F)** Animals of genotype *ieSi57[eft-3p>TIR1::mRuby]*; *ral- 1(re218re319[AID*::mKate2::3xFLAG::ral-1])* display auxin-dependent enhancement of the 2° and 3° dendritic branching defect conferred by *exoc-8*(*ok2523*). ****<0.0001, n.s. = not significant (*t-*test).

**Figure S4:**
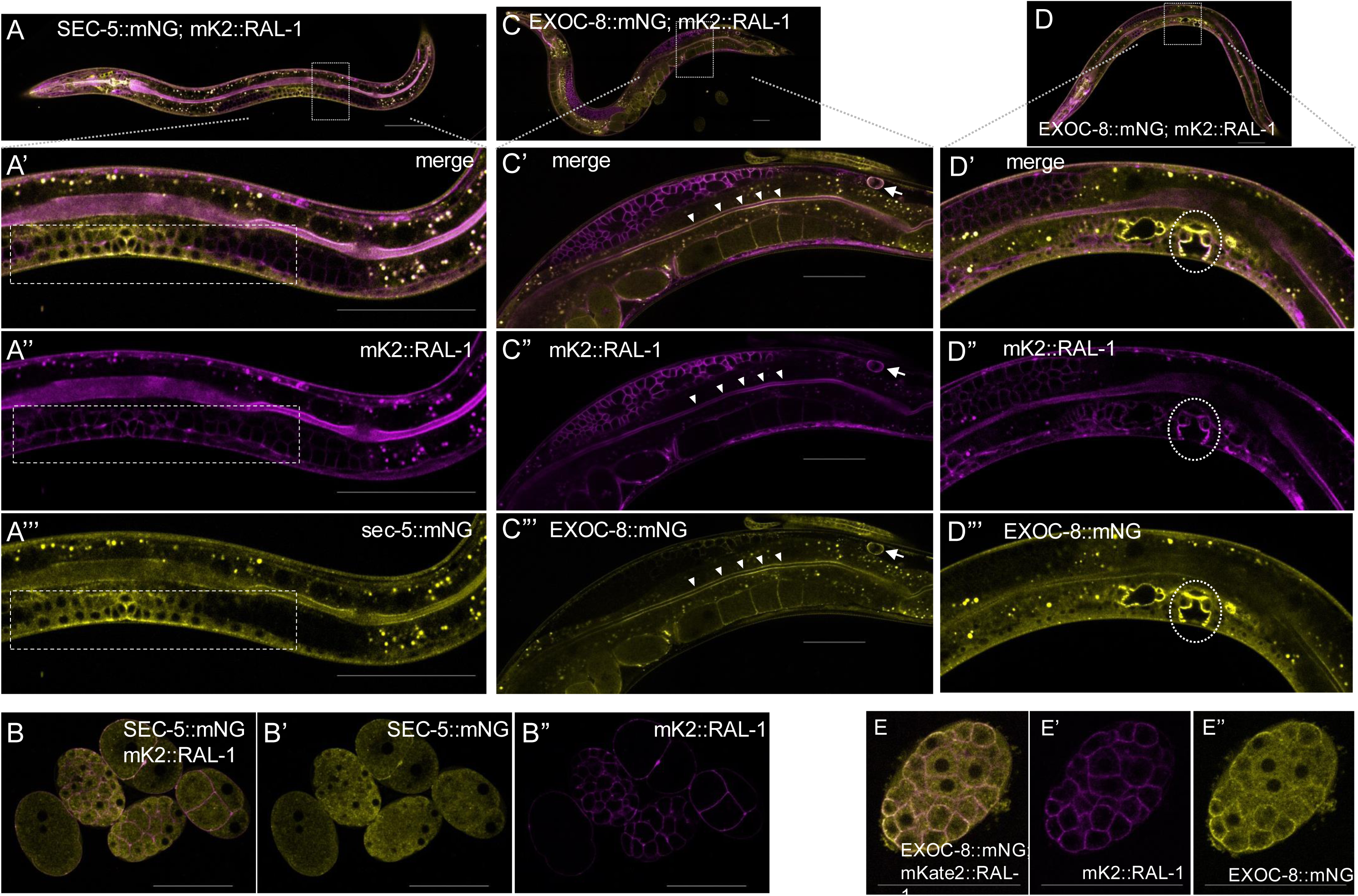
Whole-animal **(A)** and zoomed spinning disk confocal photomicrographs of merged **(A’)**, magenta **(A’’)** and yellow **(A’’’)** spinning disk confocal photomicrographs of mKate2::RAL-1 with SEC-5::mNG colocalized on the plasma membrane of VPC and vulval precursor cells (VPCs; dotted line rectangle) of a late L3 animal. **B-B’’)** The same for a group of embryos. **C-C’’’)** Whole animal **(C)** and zoomed images of of merged **(C’)** magenta **(C’’)** and yellow **(C’’’)** spinning disk confocal photomicrographs mKate2::RAL-1 with EXOC-8::mNG colocalized on the plasma membranes of a coelomocyte (arrow) and apical brush border of intestinal cells (solid triangles) of an adult animal, **D-D’’’)** on the apical membranes of invagination 22 VPC lineage progeny (dotted circle), and **E-E’’’)** on the plasma membranes of an embryo. Scale bars = 50 µm.

**Figure S9:**
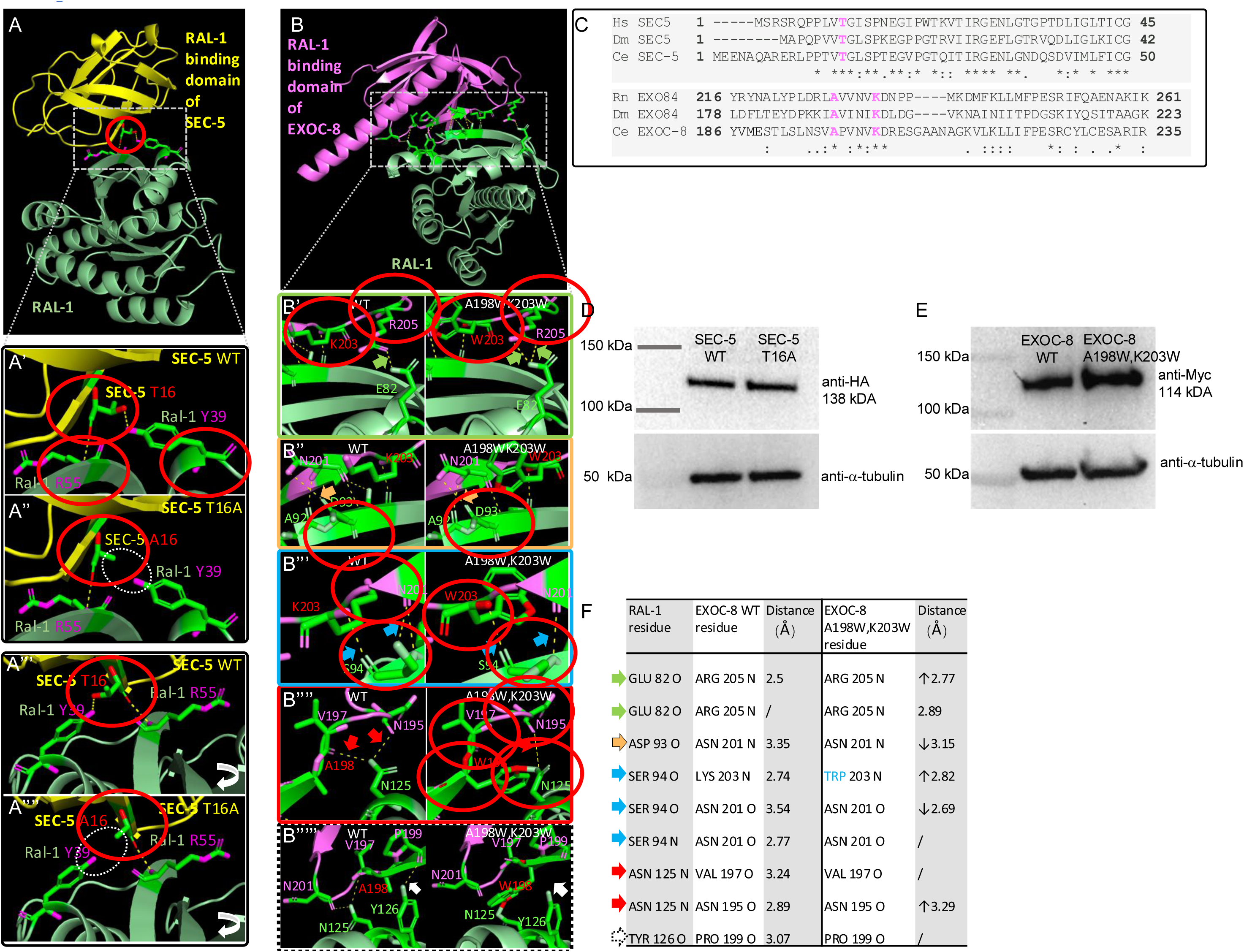
Structural models of Ral-effector interface mutations of SEC-5 and EXOC-8. **A)** Alphafold2 predicted model of SEC-5 interaction with RAL-1. With zoom-in photos **(A’)** and rotated angle **(A’’)** of WT SEC-5 (upper photos **A’** and **A’’**) and T16A mutant SEC-5 (lower photos of **A’** and **A’’**) and RAL-1 interaction domains. **B,F)** alphafold2 predicted model of EXOC-8 interaction with RAL-1. Zoom in photos are WT (**B’-B’’’’’** left photos) vs. A198W,K203W mutant EXOC-8. Color arrows pointing at changed polar contact distance, matches Table in **F**. **C)** Alignment of SEC-5 T16 flanking sequences (upper) and EXOC-8 A198, K203 flanking sequences of *C. elegans* (CE), homo sapiens (Hs) and Drosophila melanogaster **(D)**. **D,E)** T16A mutation of SEC-5 **(D)** and A198WK203W of EXOC-8 **(E)** did not destabilized protein showed by western blot.

